# Mapping tsetse fly connectivity in Uganda with machine learning landscape genetics

**DOI:** 10.64898/2026.09.11.750686

**Authors:** Norah P. Saarman, Ryan Griffiths, Anusha Bishop, Camilla Moses, Emily Calhoun, Ethan Meredith, Rosemary Bateta, Winnie A. Okeyo, Paul O. Mireji, Grace Murilla, Sylvance Okoth, Robert Opiro, Richard Echodu, Serap Aksoy, Adalgisa Caccone

## Abstract

**Introduction:** Tsetse flies (genus *Glossina*) are biting insects that transmit human and animal trypanosomiases across sub-Saharan Africa, and sustainable vector control depends on understanding dispersal barriers and reinvasion routes. Despite major progress toward elimination, Uganda remains at risk for both human forms of the disease (*Trypanosoma brucei gambiense* and *T. b. rhodesiense*) and planners still lack reliable maps of tsetse movement and reinvasion risk.

**Methods and Results:** We address this gap with machine-learning landscape genetics and species distribution models, integrating estimates of population genetic distance and geospatial environmental data to predict and map *Glossina fuscipes fuscipes* connectivity across Uganda and western Kenya. Inputs included microsatellite genotypes from 11 loci genotyped in 2,736 flies sampled from 87 localities and remotely sensed environmental predictors summarized along least-cost paths. Random forest models predicted patterns of genetic differentiation better than distance-only models, supporting the use of a machine-learning framework for connectivity inference across complex heterogeneous landscapes, and identified variables related to temperature and water availability as the strongest predictors of genetic connectivity.

**Conclusions:** Combining landscape genetics predictions of connectivity with a species distribution model revealed regions with high habitat suitability but low connectivity that represent priority zones for area-wide integrated pest management strategies, including established riverine control tools such as tiny targets and other targeted interventions aimed at reducing reinvasion risk. These results provide biologically interpretable maps and quantitative uncertainty metrics that can guide targeted tsetse control, while providing a transferable analytical pipeline for modeling and mapping genetic connectivity across other species and landscapes.

## INTRODUCTION

Tsetse flies (genus *Glossina*) are the sole cyclical vectors of trypanosomes that cause human African trypanosomiasis (HAT) and African animal trypanosomoses (AAT), diseases that continue to impose major public health and agricultural burdens across sub-Saharan Africa (Anene 2001; Baker 2013; Franco et al. 2014). Because biological transmission depends entirely on the presence and movement of tsetse populations, sustained disease control requires understanding how these vectors persist and disperse across landscapes. Uganda has historically been affected by both forms of HAT: *Trypanosoma brucei gambiense* (gHAT), responsible for a chronic form endemic in northwestern Uganda, and *T. b. rhodesiense* (rHAT), an acute zoonotic form prevalent in eastern Uganda and the Lake Victoria region. Control efforts targeting *Glossina fuscipes fuscipes*, the primary tsetse vector in Uganda, have dramatically reduced HAT incidence in recent decades and led to validation of gHAT elimination as a public health problem in Uganda. However, sustained elimination remains threatened by ongoing rHAT risk in eastern Uganda, the possibility of gHAT reintroduction into northwestern Uganda from neighboring endemic areas, and potential reinvasion of *G. f. fuscipes* populations into areas where control has successfully reduced fly densities (Franco et al. 2024; World Health Organization 2025).

Effective vector control depends not only on knowing where flies occur, but also on understanding how populations move across landscapes and recolonize areas where control has reduced fly density. Traditional trapping provides snapshots of fly presence and abundance but offers limited insight into dispersal and recolonization dynamics. These processes determine how rapidly areas targeted by control are reinfested, and have historically undermined the long-term success of tsetse control programs (Rogers and Randolph 1985; Ilemobade 2009). Population genetic measures of connectivity help address this limitation by revealing dispersal corridors, barriers, and hidden linkages among fly populations that are not detectable through trapping alone. Because genetic connectivity reflects successful dispersal and reproduction across generations, it provides a powerful framework for identifying reinvasion pathways and defining spatial units for vector control.

Environmental variation is a key factor that mediates how animal populations move across and locally adapt within landscapes, with some spatial features more important than others for connectivity. Processes influencing connectivity are of particular importance for disease vectors, where dispersal patterns can influence both the spread of pathogens and the success of control interventions. The genus *Glossina* is expected to have clear patterns of environmentally mediated dispersal because it has small effective population sizes and relatively limited dispersal over ecological timescales, both due to its unique life history as a viviparous insect (Challier, A 1982; Cuisance, D et al. 1985; Bouyer et al. 2007; Abila et al. 2008; Aksoy et al. 2013). Environmental conditions are particularly relevant in shaping patterns of connectivity for riverine species, such as *G. f. fuscipes*, where reproduction and survival are closely tied to local moisture and temperature conditions and access to shaded riparian habitat (Bursell 1958; Brightwell et al. 1992; Hargrove 2004; Krafsur 2009; Kleynhans et al. 2014). Indeed, previous population genetic studies of *G. f. fuscipes* in Uganda have revealed substantial spatial structure associated with geographic and ecological features such as lakes, river systems, and human-modified landscapes (Abila et al. 2008; Beadell et al. 2010; Hyseni et al. 2012; Echodu et al. 2013; Opiro et al. 2017; Saarman et al. 2019). These findings highlight the importance of landscape and environmental features in structuring tsetse populations and suggest that environmental drivers of dispersal could be used to predict connectivity across unsampled regions (Bouyer et al. 2015; Bouyer and Lancelot 2018; Saarman et al. 2018; Bishop et al. 2021).

The landscape genetics framework integrates landscape ecology with population genetics, most often estimates of population- or individual-level genetic distance based on allele frequency differences across the landscape (Balkenhol et al. 2009, Shirk et al. 2017, Klinga et al. 2019, Beninde et al. 2024), to test how environmental heterogeneity structures gene flow (Manel et al. 2003). Since the 2000s, complex methods have been developed to identify relationships between genetic distance and ecological predictor variables (Balkenhol et al. 2009). However, only recently have the development of machine-learning approaches such as random forest regression enabled researchers to more accurately capture nonlinear relationships and evaluate a broad predictor set rather than restricting variables *a priori*, improving inference of patterns and drivers of genetic connectivity across complex landscapes (Breiman 2001; Liaw, A and Wiener, M 2002; Prasad et al. 2006; Pless et al. 2021; Palm et al. 2023; Vanhove and Launey 2023; Day et al. 2024). Recent work applying this framework to tsetse demonstrated that integrating environmental predictors with genetic data can improve prediction of spatial patterns of connectivity relevant to vector control in *Glossina pallidipes,* a savannah species (Bishop et al. 2021). Simulation-based evaluation further supports this framework by showing that the random forest least-cost transect approach applied in Pless et al. (2021) and Bishop et al. (2021) performs substantially better than MLPE-based methods under multivariate scenarios (Vanhove and Launey 2023). Improved performance of these methods likely stems in part from the ability of machine learning to accommodate a broad suite of environmental predictors without requiring *a priori* selection of a narrowed candidate predictor set to address correlations among variables that violate parametric assumptions. This allows the relative importance of candidate predictors to be evaluated within the modeling framework rather than predetermined at the outset, making it especially promising for understanding genetic connectivity across complex landscapes.

Here, we apply and extend a machine-learning landscape genetics framework to the riverine species *G. f. fuscipes* in Uganda and the Lake Victoria region of Kenya. Our objectives were to: (1) test whether environmental predictors improve prediction of genetic connectivity relative to a distance-only model; (2) identify and spatially interpret the environmental predictors most strongly associated with genetic connectivity; and (3) integrate predicted genetic connectivity with habitat suitability to identify spatial priorities for tsetse control. We evaluated model generalizability using leave-one-point-out cross-validation and permutation tests, demonstrating substantially improved predictive performance over a distance-only baseline model. We also evaluated and optimized the spatial projection procedure by testing how alternative projection parameters affected agreement between the projected resistance surface and observed genetic connectivity, providing a direct test of the spatial output used for connectivity inference rather than relying solely on internal random forest model performance. This approach provides a scalable framework for translating landscape genetics models of connectivity into operational tools for vector control and contributes to efforts to sustain progress toward elimination of human and animal trypanosomiases.

## METHODS

### Methods Overview

We used random forest regression to predict genetic distance in *G. f. fuscipes* across Uganda and western Kenya using genetic data from 11 microsatellite loci (n = 2,736 individuals from 87 sampling sites; Figure 1). The response variable, Cavalli-Sforza and Edwards chord distance (CSE; Cavalli-Sforza and Edwards 1967), was predicted using a broad suite of 25 remotely sensed environmental predictors representing bioclimatic and physical landscape features, sampling density, and geographic distance. The workflow is presented in Figure 2 and consisted of genetic and geographic data preparation, sample filtering and data harmonization, random forest modeling, spatial projection and evaluation, leave-one-point-out cross-validation, and comparison with distance-only and permuted baseline models. Model outputs were then integrated with habitat suitability and environmental predictors to support spatial interpretation of connectivity and implications for vector control.

**Figure 1.**
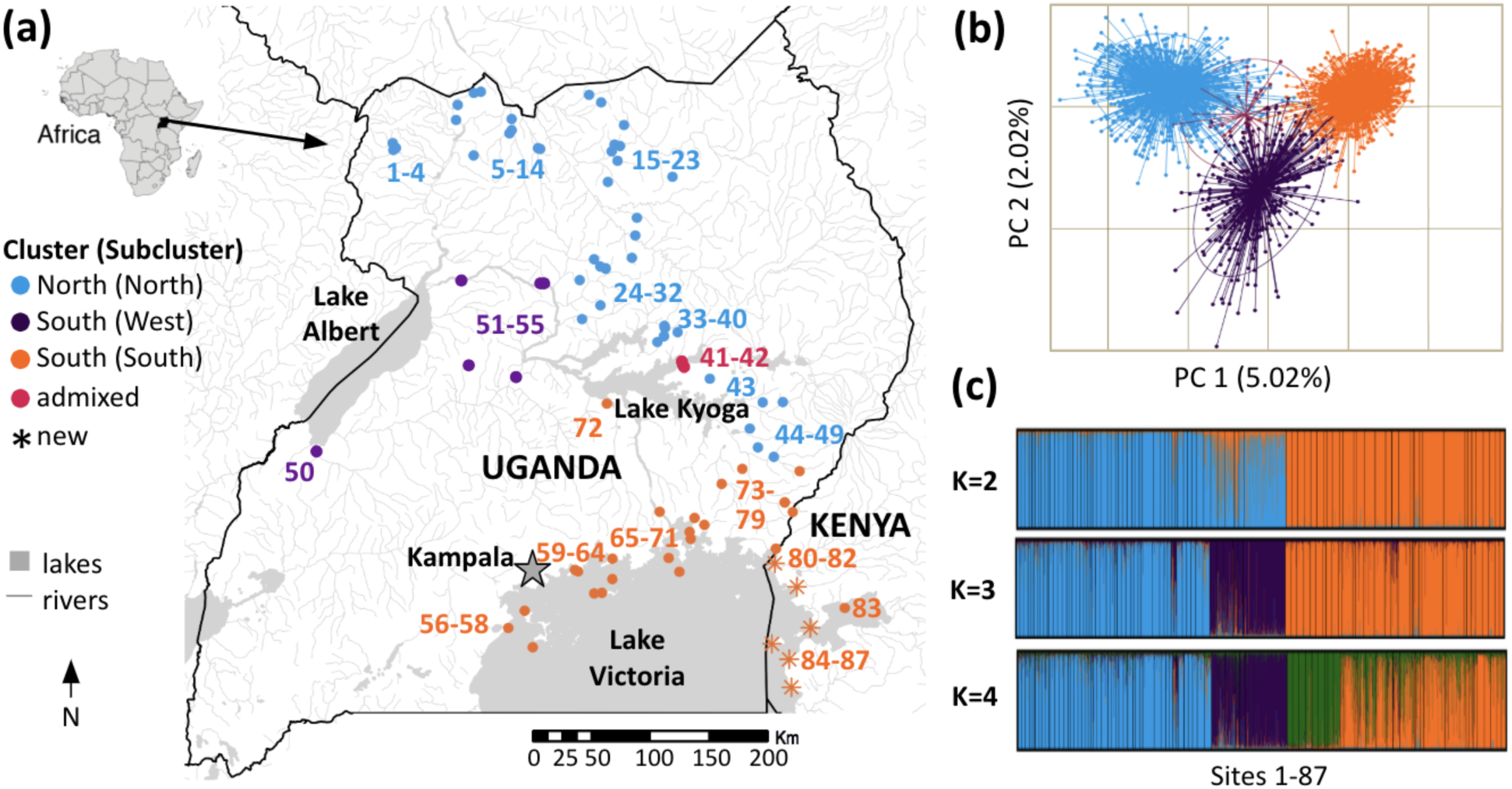
Sampling scheme and population structure of *Glossina fuscipes fuscipes* in Uganda and western Kenya. (a) Map of 87 sampling sites colored by genetic cluster assignment, with newly genotyped sites indicated by asterisks. (b) Principal Components Analysis (PCA) calculated in adegenet v2.1.4 (Jombart 2008), representing genetic diversity across 11 microsatellite loci (n = 2,736 individuals from 87 sampling sites). PCA indicates two genetic clusters, with admixed sites 41 and 42 occupying intermediate positions in PCA space. (c) STRUCTURE v2.3.4 (Pritchard et al. 2000) barplots at K = 2, K = 3, and K = 4, showing individual ancestry proportions for all sites ordered by geographic location shown in the map.

**Figure 2.**
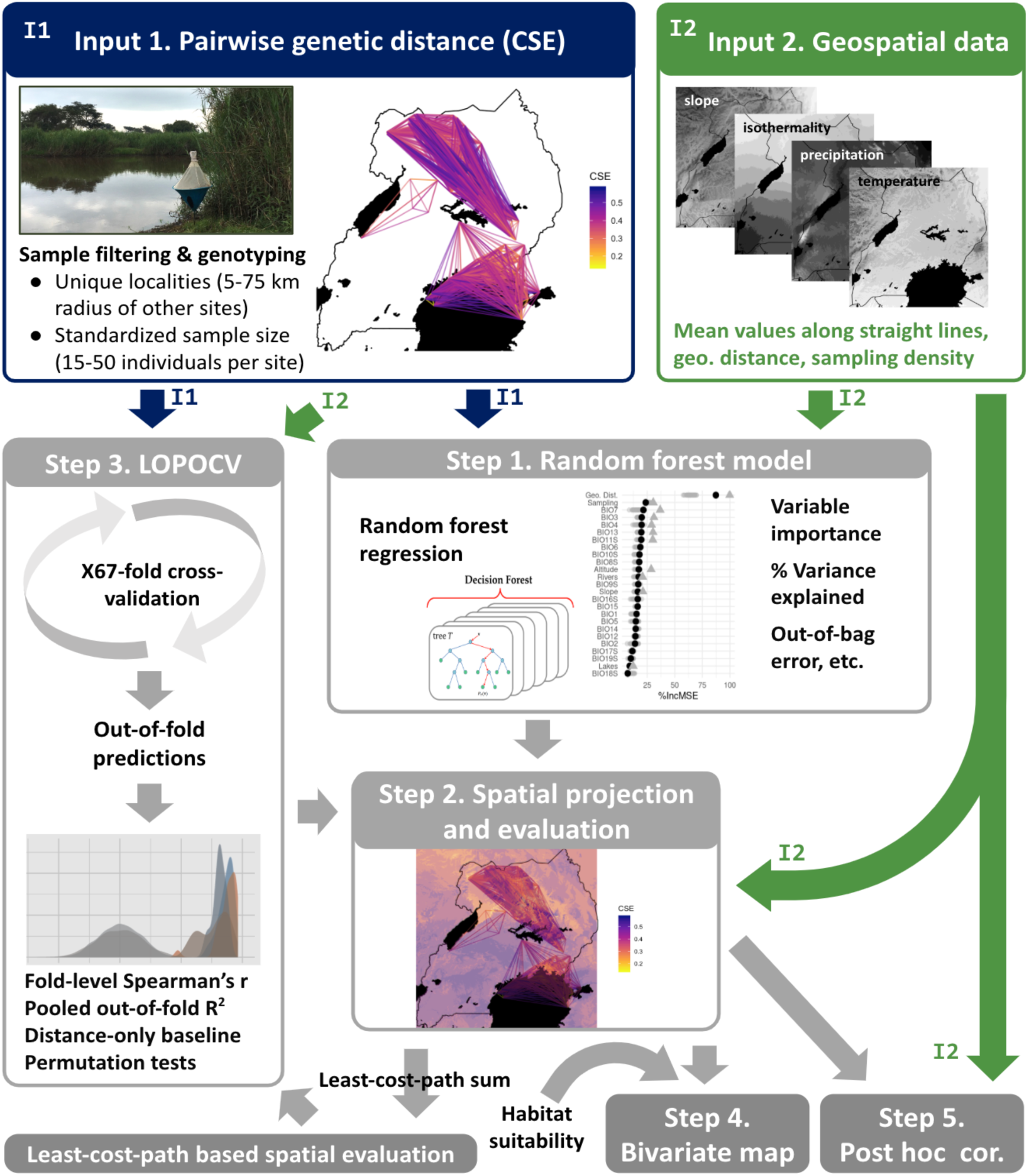
Workflow for genetic connectivity modeling. Input 1 (I1): Pairwise genetic distance, filtered for unique localities (within a 5-75 km radius of other sites) and standardized sample size (minimum of 15 individuals per site and down-sampled to a maximum of 50 individuals per site). Input 2 (I2): Geospatial environmental data. Step 1: Random forest model variable choice, performance, and variable importance. Step 2: Projected connectivity and least-cost-path-based spatial evaluation. Step 3: Leave-one-point-out cross-validation, calibration, pooled metrics, comparison with a distance-only baseline model, and permutation tests. Step 4: Bivariate map. Step 5: Post-hoc correlations with environmental predictors.

### Input 1: Tsetse Collections and Genetic Data

#### Species ecology and study region

*Glossina fuscipes fuscipes* is a member of the *Palpalis* group (subgenus *Nemorhina*) of genus *Glossina* and is associated primarily with riverine and lacustrine habitats, where it is typically restricted to shaded vegetation along riverbanks, lake shores, and gallery forest (Krafsur 2009; Aksoy et al. 2013). Populations included in this study occupy riparian and lakeside habitats throughout Uganda and western Kenya, including the Lake Victoria basin (Ford, J 1971; Pollock, JN 1982; Rogers and Randolph 1985; Jordan, AM 1993; Rogers and Robinson 2004; Cecchi et al. 2008; Ngari et al. 2020). Within this distribution, major historical genetic structure separates northern and southern populations, with additional divergence associated with the opening of the Eastern Rift Valley and the formation of the Lake Victoria basin (Saarman et al. 2019). This system is therefore well suited to landscape genetic analyses because dispersal is limited over ecological timescales and connectivity is expected to be strongly shaped by hydrologically structured habitat.

#### Genetic dataset, response variable, and sample selection

The response variable, Cavalli-Sforza and Edwards chord distances (CSE; Cavalli-Sforza and Edwards 1967), was calculated using adegenet from population genetic data comprising 11 microsatellite loci genotyped for 2,736 *G. f. fuscipes* individuals collected from 87 localities across Uganda and western Kenya between 2006 and 2018 (Figure 1). New genotyping of 166 individuals from six Kenyan sites (Supplementary Methods) was combined with previously published data from 81 localities (Beadell et al. 2010; Echodu et al. 2011, 2013; Hyseni et al. 2012; Manangwa et al. 2017; Opiro et al. 2017; Saarman et al. 2018, 2019). Population structure was evaluated using STRUCTURE v2.3.4 (Pritchard et al. 2000), with the optimal number of clusters evaluated using the Evanno method in STRUCTURE Harvester (Evanno et al. 2005; Earl and von Holdt 2012) and replicate runs summarized using CLUMPAK (Kopelman et al. 2015), and independently using principal components analysis (PCA) in adegenet v2.1.4 (Jombart 2008). Population genetic summary statistics were calculated using pegas v1.3 (Paradis 2010), hierfstat v0.5.11 (Goudet and Jombart 2022), and poppr v2.9.8 (Kamvar et al. 2015), and isolation by distance was evaluated using Mantel tests in ade4 v1.7-23 (Dray and Dufour 2007) and Mantel correlograms in vegan v2.7-1 (Oksanen et al. 2025). Detailed methods and parameters for genotyping and population genetic analyses are provided in the Supplementary Methods, with corresponding results in the Supplementary Results, Figures S1-S3, Tables S1-S3).

Because choice of genetic distance metric can influence landscape resistance inference (Beninde et al. 2024), we based our selection of CSE as the response variable on three considerations. First, CSE is a geometric measure of genetic differentiation that avoids some of the population-model assumptions associated with FST-based measures (Cavalli-Sforza and Edwards 1967). Second, previous landscape genetic studies have demonstrated the utility of CSE for estimating relative genetic distances among populations, including in the presence of missing data (Bouyer et al. 2015; Tumas et al. 2018). Third, in our previous application of this modeling framework, CSE performed better than linearized FST, although the resulting resistance surfaces were broadly similar (Pless et al. 2021).

To focus on contemporary patterns of gene flow, only pairwise CSE values between localities within the same major genetic clusters were included as response values. Population structure and isolation-by-distance analyses supported two major clusters (Supplementary Methods, Results, Figures S1-S3), with previously identified western and southern subclusters combined into a single southern cluster (Beadell et al. 2010; Echodu et al. 2011, 2013; Hyseni et al. 2012; Manangwa et al. 2017; Opiro et al. 2017; Saarman et al. 2018, 2019).

Site selection then followed a sequential filtering procedure. We retained non-admixed localities with at least 15 individuals to provide sufficient sampling for population-level estimates of genetic distance, with random down-sampling to 50 individuals to limit differences in sampling effort among localities. Where sampled localities occurred within 5 km of one another, only the locality with the largest sample size was retained. Retained sites were also required to be within 75 km of at least one other site. The 5-km thinning distance was chosen because it represents the lower bound of published estimates of tsetse dispersal of approximately 5-8 km (Challier 1982; Cuisance et al. 1985; Bouyer et al. 2007). This yielded the 67-site dataset used for downstream connectivity analyses (Table 1; Figure 2; Supplementary Table S2).

**Table 1.** Summary of sample sizes, genetic diversity statistics and pairwise Cavalli-Sforza and Edwards (1967) chord distance (CSE) across all retained sites and within the two major genetic clusters used in downstream analyses. Overall and per cluster number of retained sites (N Sites) and the number of site pairs included in the final connectivity model (N Pairs) are shown, followed by the minimum, mean, and maximum values for sample size per site (N), rarefied allelic richness (AR), Shannon-Wiener index of multilocus genotype diversity (H), multilocus linkage disequilibrium (rbarD), inbreeding coefficient (FIS), observed heterozygosity (Ho), expected heterozygosity (He), and pairwise genetic distance (CSE). Genetic diversity statistics were calculated using standard population genetic methods implemented in pegas, hierfstat, and poppr (Paradis 2010; Goudet and Jombart 2022; Kamvar et al. 2015).

| Sample | N Sites | N Pairs | Statistic | N | AR | rbarD | FIS | Ho | He | CSE |
| --- | --- | --- | --- | --- | --- | --- | --- | --- | --- | --- |
| Overall | 67 | 1091 | Minimum | 15 | 2.11 | -0.03 | -0.10 | 0.26 | 0.29 | 0.14 |
|  |  |  | Mean | 30.27 | 3.39 | 0.01 | 0.05 | 0.51 | 0.54 | 0.40 |
|  |  |  | Maximum | 50 | 4.58 | 0.07 | 0.30 | 0.69 | 0.72 | 0.68 |
| North | 35 | 595 | Minimum | 15 | 2.73 | -0.03 | -0.06 | 0.42 | 0.43 | 0.16 |
|  |  |  | Mean | 24.89 | 3.81 | 0.01 | 0.08 | 0.56 | 0.61 | 0.37 |
|  |  |  | Maximum | 50 | 4.58 | 0.07 | 0.20 | 0.69 | 0.72 | 0.57 |
| South | 32 | 496 | Minimum | 15 | 2.11 | -0.02 | -0.10 | 0.26 | 0.29 | 0.14 |
|  |  |  | Mean | 36.16 | 2.93 | 0.01 | 0.03 | 0.45 | 0.47 | 0.42 |
|  |  |  | Maximum | 50 | 4.13 | 0.05 | 0.30 | 0.63 | 0.66 | 0.68 |

### Input 2: Geospatial Data

#### Remotely-sensed environmental data

Environmental data used in this study included 1-km resolution raster layers representing 19 bioclimatic variables, slope, altitude, river density, sampling density, and a binary lake layer. These predictors were selected to represent broad climatic, hydrological, and topographic conditions relevant to the riverine and lacustrine ecology of *G. f. fuscipes* (Ford, J 1971; Pollock, JN 1982; Rogers and Randolph 1985; Jordan, AM 1993; Rogers and Robinson 2004; Cecchi et al. 2008; Ngari et al. 2020). Consistent with our machine-learning framework, we intentionally retained a broad set of biologically plausible environmental predictors rather than selecting a restricted subset *a priori*, allowing correlated predictors and nonlinear relationships to be evaluated within the model. Geographic distance and sampling density were included to account for isolation by distance and sampling structure, respectively.

The 19 bioclimatic variables were calculated from raster files downloaded from Climatologies at High Resolution for the Earth’s Land Surface Areas (CHELSA v2.1; (Karger et al. 2017) for the period 2008-2013 using the R package dismo v1.3-16 (Hijmans, RJ et al. 2024). We used seasonal bioclimatic variables based on precipitation seasonality trends observed in the study area (Wet Season 1, April–May; Dry Season 1, June–July; Wet Season 2, August– September; Dry Season 2, October–March) to better capture seasonal variation more relevant to local ecology than the default quarterly summaries. Slope and altitude raster files were obtained from the Geomorpho90m dataset (Amatulli et al. 2020) and Multi-Error-Removed Improved-Terrain (Yamazaki et al. 2017), respectively.

Sampling density and river density layers were generated using kernel density estimation in the R package *KernSmooth* v2.23-22 (Wand 2023). Sampling density was calculated from the final sampling localities included in the CSE matrix using a 20-km bandwidth, approximately corresponding to the local spacing among sampling sites (mean nearest-neighbor distance = 17.6 km). River density was calculated from river shapefiles downloaded from DIVA-GIS (March 2020). Sensitivity to river kernel bandwidth was evaluated by refitting the final model using 1-, 2-, 3-, 5-, and 10-km bandwidths while holding all other predictors and model settings constant (Supplementary Table S5).

A binary lake raster represented open water as a potential dispersal barrier, supported by previous evidence that large areas of open water can restrict *G. f. fuscipes* dispersal and gene flow (Beadell et al. 2010; Echodu et al. 2013). A uniform geographic-distance raster was included to quantify path length between site pairs. Final raster layers were clipped to the extent of Uganda and western Kenya (longitude 28.60°–35.40°, latitude −1.50°–4.73°) and projected to WGS 84 using the R package *sf* v1.0-16 (Pebesma 2018; Pebesma and Bivand 2023). Environmental conditions between site pairs were summarized by extracting raster values along least-cost paths that avoided lakes.

#### Spatial operations and visualization

All spatial operations, including raster preparation, extraction, masking, projection, and visualization of environmental inputs and model outputs, were conducted in R using raster v3.6-32 (Hijmans, RJ 2025) and ggplot2 v3.5.2 (Wickham 2009). Figures were assembled using ggpubr v0.6.1 (Kassambara 2019), ggrepel v0.9.6 (Slowikowski, 2024), and patchwork v1.3.2 (Pedersen, TL 2020).

### Connectivity Model

#### Step 1. Random forest model variable choice, performance, and variable importance

Figure 2 shows the overall workflow for genetic connectivity modeling. The response variable was pairwise CSE calculated between the retained sampling sites. To focus on contemporary patterns of gene flow, we restricted the response variable to within-cluster CSE values for the two major genetic clusters retained for modeling, with 35 sites from the North and 32 sites from the South (Table 1). We intentionally included a broad set of environmental predictors rather than using a narrow list selected from *a priori* expectations, including 19 bioclimatic variables, slope, altitude, river density, a binary lake layer, sampling density, and the sum of geographic distance along the same paths. We initially included mean, median, and mode summaries of these raster layers extracted along least-cost paths avoiding lakes between site pairs. Preliminary models showed slightly higher performance for mean summaries (85.5% variance explained) than median (84.9%) or mode (84.5%) summaries, thus we retained mean summaries for subsequent modeling.

To evaluate correlation structure among environmental predictors and aid interpretation of variable importance, we performed principal components analysis on the mean environmental predictor matrix used in the random forest model using the prcomp function from the stats package. We used the PCA to visualize major axes of covariation among predictors and to identify groups of highly correlated variables. PCA biplots and variable contribution plots were generated using the R package factoextra v2.0.0 (Kassambara and Mundt 2026), and were employed to guide selection of a reduced predictor set for the PCA-pruned comparison model. We then fit the PCA-pruned comparison model to aid interpretation of variable importance while retaining the full predictor set in the final model.

We modeled environmental predictors of genetic connectivity using random forest regression, generally following Bishop et al. (2021) and Pless et al. (2021). Analyses were conducted in R with the randomForest package v4.7-1.2 (Liaw and Wiener 2002). The final model used the full set of mean-based predictors. We optimized mtry, the number of predictor variables considered at each split, using tuneRF, with 500 trees per candidate model, a step factor of 1.5, and a minimum relative improvement in OOB error of 0.01. The search selected mtry = 12 for the final model. Initial model performance was evaluated using the out-of-bag resampling procedure in randomForest, which estimates percent variance explained and mean squared error (MSE). Predictor importance was ranked primarily by percent increase in mean squared error (%IncMSE), with increase in node purity examined for comparison. Variable-importance rankings from leave-one-point-out cross-validation models and the PCA-pruned comparison model were compared to assess stability across modeling formulations.

#### Step 2. Projected connectivity and least-cost-path-based spatial evaluation

The final fitted model was projected across Uganda and western Kenya using the same environmental raster stack used to fit the model. For the least-cost-path-based spatial evaluation, predicted CSE values were used as resistance surfaces, with lake cells assigned high cost equal to the maximum value of the projected surface, reflecting evidence that stretches of open water greater than 10 km can act as barriers to *G. f. fuscipes* dispersal and gene flow (Beadell et al. 2010; Echodu et al. 2013; Saarman et al. 2018). Sensitivity to this assumption was evaluated using lake resistance values of 0.5, 1, 1.5, and 2 times the maximum predicted terrestrial resistance. Least-cost paths were then recalculated among site pairs. For each projected surface, predicted CSE values were summarized as the sum of values along least-cost paths (LCP_sum), calibrated to observed CSE by linear regression, and evaluated using Spearman correlation, coefficient of determination (R2), mean squared error (MSE), root mean squared error (RMSE), and mean absolute error (MAE).

Because geographic distance was included as a predictor, we screened alternative fixed projection distances for the least-cost-path-based spatial evaluation step by substituting constant path-length values into the projection stack and comparing candidate values from 1 to 100 km. Based on this screening, we carried forward a projection distance of 1 km for downstream spatial evaluation, iterative updating, and visualization. We also explored iterative updating by refitting the random forest model and recalculating least-cost paths across 10 iterations, evaluating each iteration using the same internal and spatial performance metrics.

#### Step 3. Leave-one-point-out cross-validation, calibration, and permutation tests LOPOCV random forest model performance

To further evaluate model generalizability, we implemented leave-one-point-out cross-validation (LOPOCV), in which each fold held out one sampling site. All pairwise CSE values involving that site were removed from the training dataset and used as the test set. For each fold, a new random forest model was trained on the remaining data using the same predictor set. Model performance was first summarized using random forest out-of-bag metrics, including percent variance explained and mean squared error. We then fit a calibration regression using training pairs only, with observed CSE regressed on raw predicted CSE. The resulting fold-specific calibration equation was retained to assess calibration stability across training sets and applied to the held-out pairwise predictions involving the excluded sampling site, without using the held-out observed CSE values for calibration. Fold-level predictive performance was calculated for the held-out pairs using Spearman correlation, RMSE, and MAE. Because several held-out folds had limited variance in observed CSE, fold-specific R2 values were unstable and were not emphasized. Instead, pooled out-of-fold R2 was calculated across all folds as a more stable overall measure of predictive performance.

To benchmark model performance, we repeated the LOPOCV procedure with a distance-only baseline model that included only geographic distance as the main predictor, with sampling density retained for consistency across model formulations. Observed fold-level Spearman correlations from the full random forest and distance-only baseline models were compared using paired sign-permutation tests with 10,000 permutations. We also benchmarked performance against a null expectation by repeating the full LOPOCV procedure on 100 permuted datasets in which pairwise CSE response values were randomly shuffled while the environmental predictor matrix and LOPOCV fold structure were held fixed.

#### LOPOCV least-cost-path-based spatial evaluation of projected connectivity

We then applied the same least-cost-path-based spatial evaluation framework described above to each fold-specific LOPOCV model, using a fixed projection distance of 1 km to match the best-performing method from preliminary screening. Fold-specific least-cost-path predictions were calibrated using training pairs only and evaluated on held-out pairs using the same fold-level and pooled out-of-fold metrics described above. Calibration slopes and intercepts from this spatial evaluation were also retained to assess calibration stability across folds. We also completed the LOPOCV least-cost-path-based spatial evaluation for the distance-only baseline model.

### Linking Connectivity with Environment and Habitat Suitability

#### Step 4. Bivariate map

We compared the projected CSE surface with an updated habitat suitability surface based on Wint and Rogers (2000) and refined in Saarman et al. (2018), accessed via Dryad. Suitability values below 0.05 were masked to exclude areas predicted to be unsuitable. The projected CSE surface was scaled from 0 to 1 and inverted as 1 minus scaled CSE to produce a continuous surface of inferred genetic connectivity, while the habitat suitability surface was rescaled from 0 to 1. Both surfaces were classified for visualization and plotted together as a bivariate map. The bivariate map was used to identify areas where connectivity and suitability were concordant or divergent in ways potentially relevant to area-wide integrated pest management.

#### Step 5. Post-hoc correlations with environmental predictors

For post hoc spatial interpretation, we selected five environmental predictors that ranked highly in random forest variable importance while representing relatively distinct environmental gradients: isothermality (BIO3), temperature seasonality (BIO4), temperature annual range (BIO7), mean temperature of the coldest season (BIO11S), and precipitation of the wettest month (BIO13). The full random forest model intentionally retained correlated predictors because predictor independence is not an assumption of random forest regression, allowing us to retain the broad set of biologically plausible candidate predictors rather than preselecting a restricted subset. However, correlation among predictors can complicate interpretation of their individual importance and effects. We therefore considered predictor correlation when selecting variables for post hoc analysis, choosing high-importance predictors that represented relatively nonredundant environmental gradients. For each predictor, we calculated local Pearson correlations between the predictor raster and the predicted connectivity surface using corLocal in the R package raster, with a 21-pixel sliding window. This allowed the direction and strength of predictor-response relationships to vary spatially across the study region.

## RESULTS

### Input 1: Population Genetic Analysis

Population genetic analyses supported two major genetic clusters, North and South (Figure 1), with the six newly genotyped western Kenya sites assigned to the South cluster (Supplementary Results, Tables S1-S3, Figures S1-S3). After excluding admixed sites 41 and 42, geographically isolated site 50, sites with fewer than 15 individuals, and redundant nearby samples from overlapping trapping localities, the final connectivity modeling dataset included 67 sites: 35 in the North cluster and 32 in the South cluster (Table 1; Supplementary Table S2). Pairwise CSE values averaged 0.40 across all retained site pairs and were slightly lower in the North than in the South (0.37 vs. 0.42; Supplementary Results, Table S3).

### Connectivity Model

#### Step 1. Random forest model performance and variable importance

The final genetic connectivity model used within-cluster pairwise CSE as the response variable and the complete mean-based predictor set after comparison with median- and mode-based summaries. The final tuned random forest model performed strongly, explaining 85.7 percent of the variance in pairwise CSE with an out-of-bag mean square error (MSE), a standard measure of model performance, of 0.0012. Model-development comparisons supported use of raw CSE above other candidate response formulations (Supplementary Table S4), while model performance was nearly identical across river-density bandwidths of 1–10 km, supporting the 3-km bandwidth used in the final model (Supplementary Table S5). Random forest out-of-bag MSEstabilized well before 500 trees (Supplementary Figure S4), supporting the use of 500 trees in the final model. The optimized mtry was 12.

Variable-importance rankings based on %IncMSE were broadly stable across the full model, the PCA-pruned comparison model, and the LOPOCV folds (Figure 3). PCA of the mean environmental predictors, shown in the inset of Figure 3, highlighted broad covariance among temperature, precipitation, and habitat-related variables. The PCA-pruned comparison model performed similarly to the full model, explaining 85.48% of the variance, and produced broadly similar variable-importance patterns. The full predictor set was therefore retained as the final model, while the PCA-pruned model provided a comparison for evaluating the robustness of variable-importance patterns to predictor covariance.

**Figure 3.**
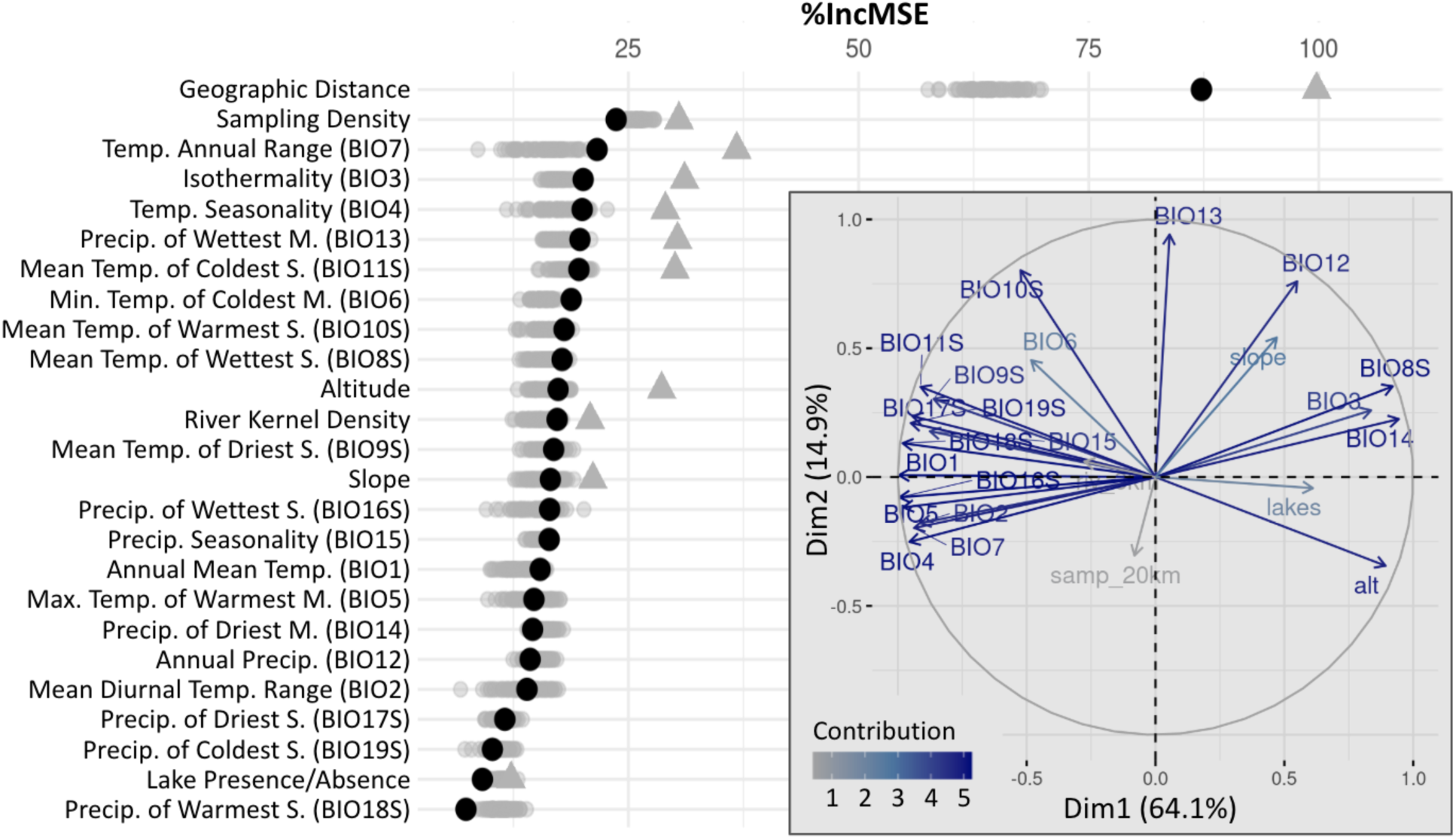
Final model variable importance (%IncMSE) and PCA of environmental predictors. Black circles show variable importance for the full model trained on 1,091 pairwise genetic distances among 67 sampling sites using all 25 predictors. Gray triangles the PCA-pruned model, trained on the same 1,091 pairwise genetic distances using a reduced set of 11 predictors and displayed to aid interpretation. Gray points show 67 LOPOCV models, each trained on pairwise genetic distances among 66 sites and evaluated against pairs involving the held-out site. The inset shows PCA structure among environmental predictors. Abbreviations: Temp. = temperature; Precip. = precipitation; Min. = minimum; Max. = maximum; S. = season; M. = month.

Geographic distance ranked first and sampling density ranked second. Among environmental predictors, temperature annual range (BIO7) was the strongest predictor, followed by isothermality (BIO3), temperature seasonality (BIO4), precipitation of the wettest month (BIO13), and mean temperature of the coldest season (BIO11S). Their importance distributions overlapped across LOPOCV folds, indicating some variation in their relative rankings among models rather than a strict ordering of importance. Additional correlated environmental predictors also ranked highly in the full model (Figure 3).

#### Step 2. Projected connectivity and least-cost-path-based spatial evaluation

Figure 4 shows inferred genetic connectivity from the final tuned random forest model, projected using a fixed projection distance of 1 km, which was selected because it performed best in spatial screening. Projected surfaces using 1, 2, and 5 km projection distances performed similarly, whereas larger projection distances progressively reduced fit, with modest declines in Spearman correlation and R^2^ and increases in error metrics (Supplementary Figure S5). Spatial performance was substantially lower when lake resistance was reduced to 0.5 times the maximum terrestrial resistance, whereas increasing lake resistance to 1.5 or 2 times the maximum produced only modest improvements over the value of 1 used in the final analysis (Supplementary Figure S5). LCP_sum was the highest performing of candidate methods for map projections (Supplementary Table S6), and the final model performed well in spatial evaluation after projection, with Spearman’s r = 0.7848, R2 = 0.6159, MSE = 0.00311, RMSE = 0.0558, and MAE = 0.0454 (Supplementary Table S6). Exploratory iterative updating of least-cost paths across 10 iterations produced only minor fluctuations in OOB model performance after the initial adjustment, with no consistent improvement in variance explained or MSE (Supplementary Figure S6).

**Figure 4.**
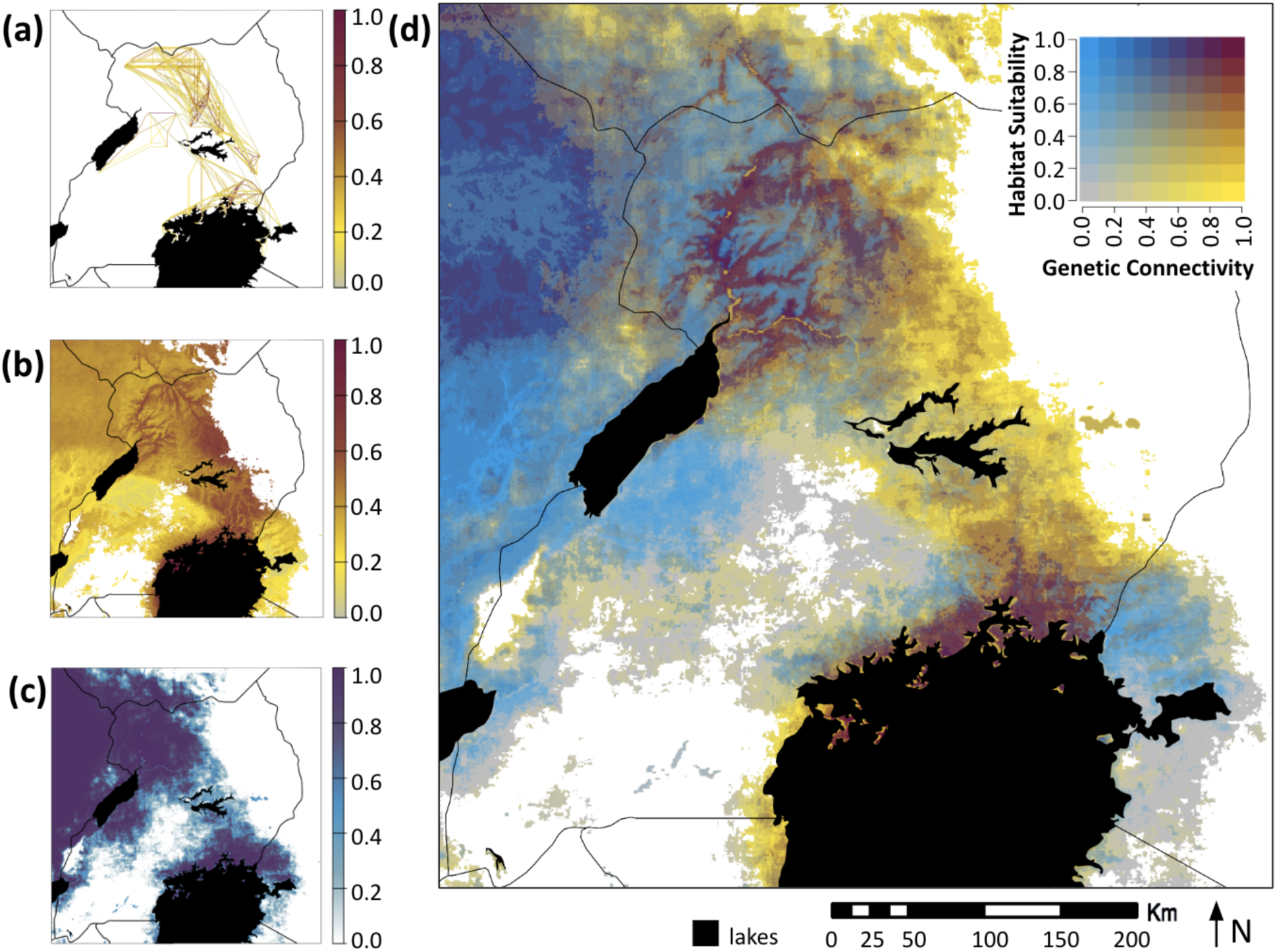
Projections of predicted genetic connectivity and habitat suitability and bivariate map. (a) Observed pairwise genetic connectivity (1 − scaled CSE) along least-cost paths between sampled sites. (b) Predicted genetic connectivity from the final random forest model (1 − scaled predicted CSE). (c) Predicted habitat suitability, combining the 2018 update and FAO distribution surface, rescaled 0–1. (d) Bivariate map of genetic connectivity and habitat suitability; dark red indicates high values of both, yellow high connectivity/low suitability, blue low connectivity/high suitability, and gray low values of both. White areas in (b–d) indicate habitat suitability <0.05 and were masked.

#### Step 3. Leave-one-point-out cross-validation, calibration, and permutation tests LOPOCV random forest model performance

Figure 5 summarizes LOPOCV performance of the full genetic connectivity model and compares it with the distance-only baseline model. Fold-specific calibration regressions for the raw model predictions were highly consistent across LOPOCV folds, indicating a stable relationship across training sets. Calibration slopes ranged from 1.056 to 1.064 (SD = 0.0016), and intercepts ranged from −0.02535 to −0.02249 (SD = 0.00062). The full model showed strong predictive performance across held-out sites, with pooled out-of-fold R2 = 0.8041, Spearman correlation = 0.8904, RMSE = 0.03985, and MAE = 0.03062. Fold-level Spearman correlations ranged from 0.4726 to 0.9629.

**Figure 5.**
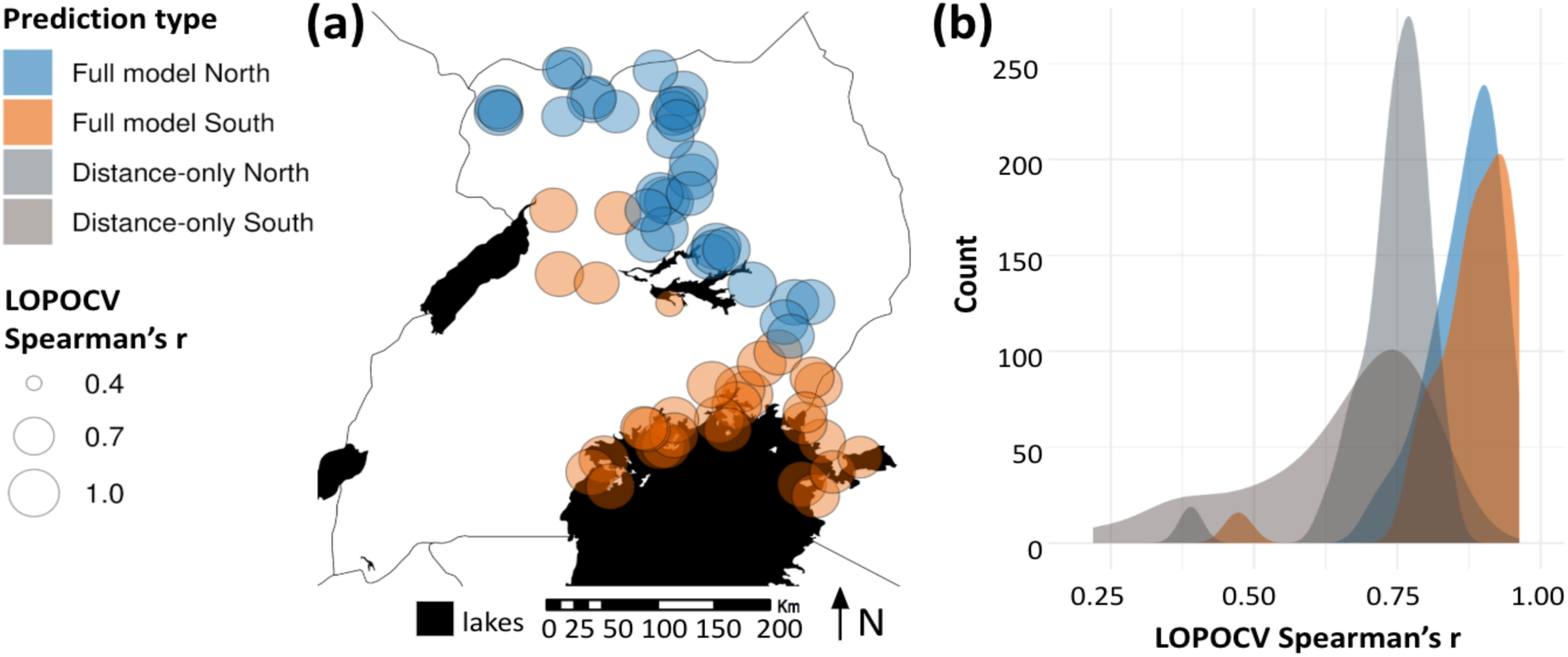
Leave-one-point-out cross-validation (LOPOCV) performance of the full genetic connectivity model versus a distance-only baseline. LOPOCV included 67 folds, each trained on pairwise genetic distances among 66 sites and evaluated against pairs involving the held-out site. Full model results are shown in blue for the North and orange for the South; distance-only results are shown in shades of gray. (a) Spatial distribution of full model LOPOCV performance, with point size proportional to Spearman’s r. (b) Distributions of fold-level Spearman’s r for the full and distance-only models, with mean values of 0.87 versus 0.74 in the North and 0.88 versus 0.65 in the South, corresponding to improvements of 0.13 and 0.23, respectively. Overall, the full model outperformed the distance-only baseline (mean difference in Spearman’s r = 0.18; paired sign-permutation test, 10,000 permutations, p < 0.0001).

The full model also outperformed the distance-only baseline, especially in the South. Mean fold-level Spearman correlations were 0.87 versus 0.74 in the North and 0.88 versus 0.65 in the South, corresponding to improvements of 0.13 and 0.23, respectively (Figure 5). Across all folds, the mean improvement in Spearman correlation was 0.18 (paired empirical sign-permutation test, 10,000 permutations, p < 0.0001), supporting the strong predictive value of the environmental predictors. Performance was also significantly higher than null expectations based on LOPOCV repeated across 100 permuted datasets, in which pairwise CSE values were randomly shuffled while the predictor matrix and fold structure were held constant (Supplementary Figure S7).

#### LOPOCV least-cost-path-based spatial evaluation of projected connectivity

Least-cost-path-based spatial evaluation of the fold-specific full models also showed strong predictive performance across held-out sites after projection to the spatial connectivity surface. Pooled out-of-fold performance was R2 = 0.5782, Spearman correlation = 0.7850, RMSE = 0.05848, and MAE = 0.04740. Fold-level Spearman correlations ranged from 0.5512 to 0.9149 (Supplementary Figure S7), indicating consistently positive predictive performance across held-out sites.

### Linking Genetic Connectivity with Environment and Habitat Suitability

#### Step 4. Bivariate map

Rescaling and inverting the projected CSE surface produced a continuous surface of inferred genetic connectivity, rather than predicted CSE, across Uganda and western Kenya (Figure 4b). Comparison with the external habitat suitability surface also scaled from 0 to 1 across the study area (Figure 4c), produced a bivariate map (Figure 4d) that showed that connectivity and suitability were not spatially equivalent, with marked areas of concordance and divergence around Lake Albert, Lake Kyoga, and the Lake Victoria basin.

#### Step 5. Post-hoc correlations with environmental predictors

Figure 6 shows three representative predictors with distinct spatial patterns, isothermality (BIO3), temperature seasonality (BIO4), and precipitation of the wettest month (BIO13), together with their local Pearson correlations with predicted connectivity. The full set of five focal predictors is shown in Figure S6. These results indicate substantial spatial heterogeneity in the direction and strength of predictor-response relationships across the landscape, with three general patterns and an approximate north-south transition. Connectivity and isothermality (BIO3) were generally positively associated near Lake Victoria but negatively associated north of Lake Kyoga and across parts of central Uganda. Connectivity was generally positively associated with temperature seasonality (BIO4) and temperature annual range (BIO7) in high-suitability areas, except along the western shore of Lake Victoria. In contrast, connectivity was generally negatively associated with precipitation of the wettest month (BIO13) and mean temperature of the coldest season (BIO11S) in the North but positively associated with these predictors in the South (Figures 6, S6).

**Figure 6.**
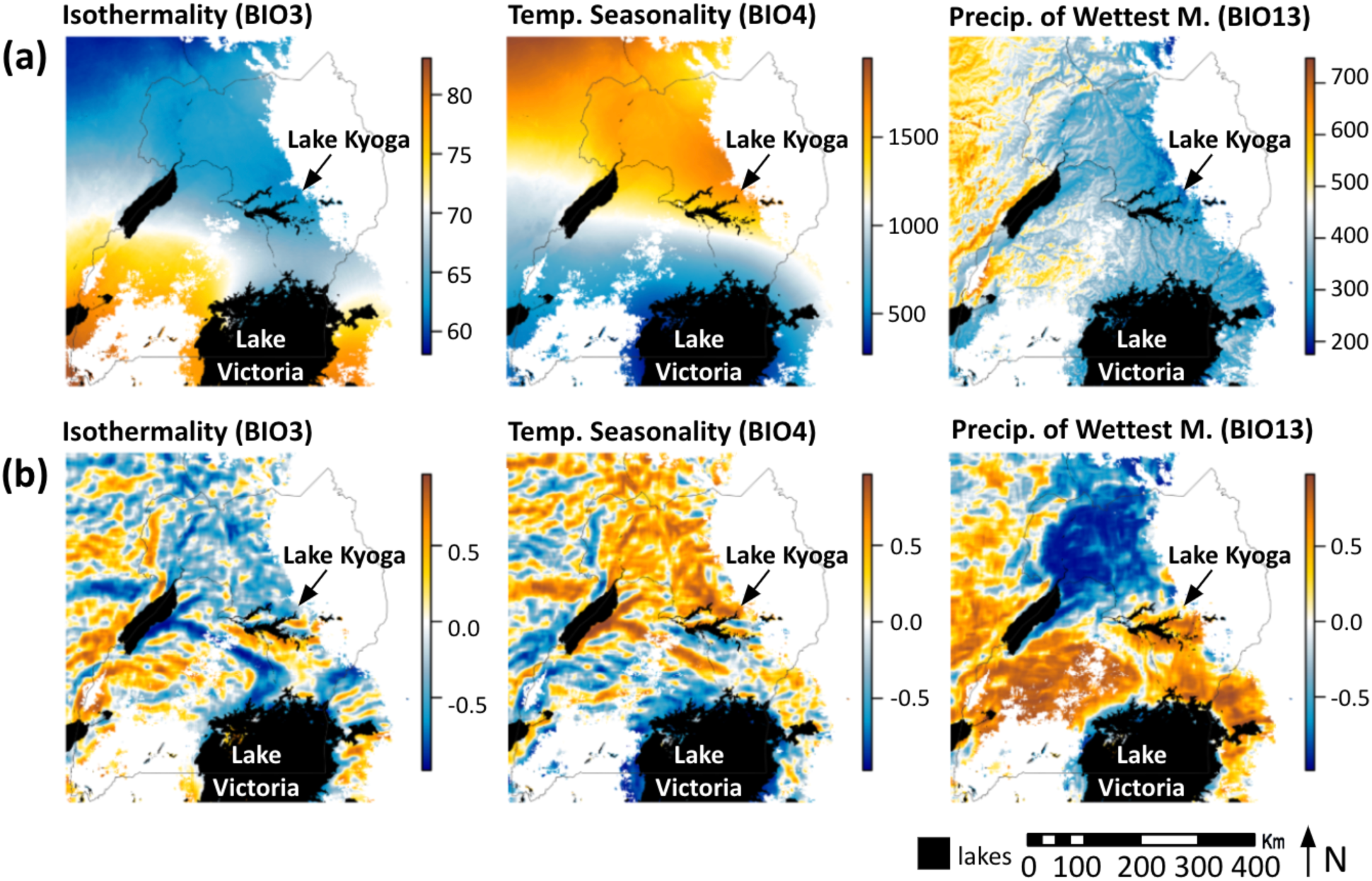
Post hoc spatial interpretation of three high-importance environmental predictors representing distinct spatial patterns. (a) Raw predictor values for isothermality (BIO3; mean diurnal range / annual temperature range x 100), temperature seasonality (BIO4; standard deviation of monthly temperature x 100, degrees C), and precipitation of the wettest month (BIO13; mm). (b) Local Pearson correlations between each predictor and predicted genetic connectivity, calculated using a 21-pixel sliding window. Positive correlations (orange) indicate areas where higher predictor values are associated with higher predicted connectivity, whereas negative correlations (blue) indicate areas where higher predictor values are associated with lower predicted connectivity. Lakes are shown in black, and white areas indicate habitat suitability <0.05 and were masked. Lake Victoria and Lake Kyoga are labeled for geographic reference.

## DISCUSSION

The goals of this study were to test whether environmental predictors improve prediction of genetic connectivity relative to a distance-only model for *G. f. fuscipes* across Uganda and western Kenya, identify and spatially interpret the environmental predictors most strongly associated with genetic connectivity, and compare predicted connectivity with modeled habitat suitability to identify spatial priorities for vector surveillance and control. Population genetic analyses identified two major genetic clusters, North and South (Figure 1), with differences in genetic diversity and differentiation summarized in Table 1. These data formed the basis for the machine-learning landscape genetics workflow used to model genetic connectivity (Figure 2). The final random forest model predicted pairwise genetic differentiation well across out-of-bag evaluation, least-cost-path-based spatial evaluation, and leave-one-point-out cross-validation, and outperformed a distance-only baseline and permuted dataset (Figure 5). Geographic distance and sampling density ranked highest overall, while temperature- and precipitation-related variables were among the most important environmental predictors (Figure 3). A bivariate map (Figure 4) comparing predicted connectivity with habitat suitability identified broad regions differing in reinvasion risk and control feasibility. Post hoc spatial analyses identified geographic variation in associations between genetic connectivity and environmental predictors related to temperature, water availability, and hydrologically structured habitat (Figure 6). For vector control, our integrative approach can help distinguish potential corridors, barriers, and relatively isolated populations, providing spatial information directly relevant to surveillance, reinvasion risk, and area-wide integrated pest management.

### Population Genetics Analysis

The analyses supported two major genetic clusters, North and South (Figure 1), broadly consistent with previous population genetic and genomic studies of *G. f. fuscipes* in Uganda. Earlier work identified a major historical division between northern and southern lineages, together with finer-scale subdivision and admixture within and between these regions (Beadell et al. 2010; Echodu et al. 2013; Opiro et al. 2017; Saarman et al. 2019). These broad-scale patterns have been linked to historical changes in drainage connectivity associated with formation of the East African Rift system and reorganization of the Lake Victoria and Nile drainage networks (Saarman et al. 2019).

Patterns of genetic diversity indicated that northern sites had higher overall diversity than southern sites, with greater mean rarefied allelic richness (3.81 vs. 2.93) and expected heterozygosity (0.61 vs. 0.47; Table 1). Pairwise CSE was also slightly higher on average in the South than in the North (0.42 vs. 0.37; Table 1), consistent with stronger genetic differentiation among southern sites. This is broadly consistent with previous evidence of restricted gene flow and substantial population structure within southern Uganda (Beadell et al. 2010; Echodu et al. 2013).

Strong positive isolation by distance in both genetic clusters further supports geographic distance as a primary constraint on gene flow in this system (Supplementary Results, Figures S4-S5), consistent with previous studies reporting significant isolation by distance and ongoing gene flow among neighboring *G. f. fuscipes* populations in Uganda (Beadell et al. 2010; Opiro et al. 2017; Saarman et al. 2019). Overall, the fine-grained differences between the North and South, together with improved performance of the full model over the distance-only baseline, indicate that environmental heterogeneity contributes additional structure to genetic connectivity beyond that explained by geographic distance alone.

### Connectivity Model

#### Step 1. Random forest model performance and variable importance

The final model explained 85.7% of the variance in pairwise CSE based on random forest out-of-bag predictions, which compares favorably with performance reported in many landscape genetic analyses (Murphy et al. 2010; Hether and Hoffman 2012; Marrotte et al. 2014; Van Strien et al. 2014; Bouyer et al. 2015; Shirk et al. 2017; Bishop et al. 2021; Pless et al. 2021; Palm et al. 2023; Vanhove and Launey 2023; Day et al. 2024). Geographic distance ranked first in variable importance, consistent with the strong isolation-by-distance pattern and historical population structure described above, while sampling density ranked second. The importance of geographic distance does not diminish the contribution of environmental predictors, as the full model substantially outperformed the distance-only baseline in LOPOCV (Figure 5), demonstrating that environmental heterogeneity explained additional variation in genetic connectivity beyond geographic distance alone.

Among environmental predictors, temperature annual range (BIO7) ranked highest, followed by isothermality (BIO3), temperature seasonality (BIO4), precipitation of the wettest month (BIO13), and mean temperature of the coldest season (BIO11S; Figure 3). Variable-importance rankings were broadly stable across the full model, PCA-pruned comparison model, and LOPOCV folds (Figure 3). However, correlation among predictors can complicate interpretation of their individual random forest variable importance values (Strobl et al. 2008; Gregorutti et al. 2017).

The importance of temperature- and moisture-related predictors is broadly consistent with tsetse biology. Temperature extremes and desiccation stress are known to influence pupal survival and the fat reserves of newly emerged teneral flies, with downstream effects on survival and reproductive potential (Bursell 1958; Brightwell et al. 1992; Hargrove 2004; Kleynhans et al. 2014). Precipitation of the wettest month may also reflect the availability of humid microclimates needed for larval deposition and puparial development (Bursell 1958; Brightwell et al. 1992; Hargrove 2004; Aksoy et al. 2013; Kleynhans et al. 2014). More broadly, these predictors are consistent with previous work showing that lakes, river systems, and hydrologically structured habitat contribute to population genetic structure in *G. f. fuscipes* (Abila et al. 2008; Beadell et al. 2010; Hyseni et al. 2012; Echodu et al. 2013; Opiro et al. 2017; Saarman et al. 2018, 2019; Bishop et al. 2021).

Comparison with the earlier application of this framework to the savannah tsetse species *G. pallidipes* suggests that species life history and ecology can strongly influence the predictability of genetic connectivity. The *G. pallidipes* connectivity model explained 67% of the variance in genetic distance based on random forest out-of-bag predictions, compared with 85.7% here, although differences in sampling design, geographic extent, and model implementation limit direct comparison of model performance (Bishop et al. 2021). In both species, climatic variables contributed substantially to predicted connectivity, but the strongest environmental associations differed. Precipitation of the driest season was the most important environmental predictor of *G. pallidipes* connectivity, whereas temperature annual range, isothermality, and temperature seasonality were among the strongest environmental predictors for *G. f. fuscipes*. These differences are consistent with the distinct ecologies of the two species, with *G. pallidipes* occupying seasonally variable savannah habitat and *G. f. fuscipes* more closely associated with riverine and lacustrine habitat. The resulting management implications also differ in spatial scale: Bishop et al. (2021) supported managing much of eastern Kenya as a single connected *G. pallidipes* unit, whereas the more spatially heterogeneous connectivity predicted here identifies finer-scale differences in potential reinvasion corridors and relatively isolated *G. f. fuscipes* populations.

#### Step 2. Projected connectivity and least-cost-path-based spatial evaluation

Least-cost-path-based spatial evaluation using LCP_sum performed well, indicating that the projected surface preserved biologically meaningful spatial relationships among sampled sites. Exploratory screening of projection distance further showed that projections using fixed distances of 1, 2, and 5 km performed similarly (Supplementary Figure S5), whereas larger projection distances reduced fit. This pattern suggests that connectivity in this system was best captured at relatively fine spatial scales.

Exploratory iterative updating of least-cost paths did not improve performance across ten iterations (Supplementary Figure S6). Instead, paths based on the initial connectivity surface, with routes constrained to avoid large lakes, performed as well as or better than subsequent paths recalculated from iteratively updated connectivity surfaces. In the mosquito species *Aedes aegypti*, iterative least-cost-path updating improved spatial prediction at the continental scale across the southern United States in the original implementation of this framework (Pless et al. 2021). In contrast, *G. f. fuscipes* is a riverine tsetse closely associated with waterways and riparian habitat. Thus, dispersal may be more strongly constrained to relatively well-defined and connected riverine habitat networks, reducing the additional information gained by iteratively refining least-cost paths. The relatively dense sampling across much of the suitable habitat in the present study may also have contributed to the stability of the initial connectivity surface. These differences emphasize that the value of iterative path updating is likely to depend on focal species biology, the scale of sampling and modeling, and landscape structure, and should therefore be evaluated rather than assumed when applying this framework to new systems.

#### Step 3. Leave-one-point-out cross-validation, calibration, and permutation tests LOPOCV random forest model performance

LOPOCV supported strong generalizability across held-out sites and showed that environmental predictors improved prediction beyond geographic distance alone (Figure 5; Supplementary Figure S7). This improvement was evident in both genetic clusters but was greater in the South, suggesting that environmental variation may play a larger role in structuring connectivity in the southern portion of the study area. Performance substantially exceeded the null expectation from permuted datasets (Supplementary Figure S7c), further supporting the predictive value of the environmental model. The lowest-performing fold, Junda (site 72 on the south shore of Lake Kyoga; Figure 5), may reflect its proximity to the major genetic transition and admixture zone identified here (Figure 1) and in previous studies (Beadell et al. 2010; Echodu et al. 2013; Saarman et al. 2019), which may not be fully captured by the broader model.

### LOPOCV least-cost-path-based spatial evaluation of projected connectivity

Least-cost-path-based spatial evaluation similarly supported generalizability across held-out sites, indicating that predictive signal was retained when the fold-specific models were projected to spatial connectivity surfaces. There was no obvious low-performing outlier in the spatial evaluation (Supplementary Figure S7b), as there was in the internal LOPOCV evaluation (Figure 5; Supplementary Figure S7a). Instead, performance was generally lower across sites in the western subcluster, where sampling was much sparser (only five sites; Supplementary Table S2), suggesting that spatial projection may be somewhat less accurate in sparsely sampled regions.

### Linking Genetic Connectivity with Environment and Habitat Suitability

#### Step 4. Bivariate map

In the bivariate map, high suitability did not always correspond to high connectivity, indicating that areas suitable for *G. f. fuscipes* persistence may differ from areas that facilitate movement and reinvasion (Figure 4). This distinction supports the conclusion that habitat suitability is not synonymous with permeability for dispersal (Mateo-Sanchez et al. 2015). Although habitat suitability surfaces are sometimes used to parameterize landscape genetic models (Milanesi et al. 2015), and functional connectivity can be incorporated into habitat suitability models (Bouyer et al. 2018), our results support keeping these outputs conceptually distinct. Habitat suitability identifies where flies may occur, whereas genetic connectivity helps identify whether populations are likely to be linked, isolated, or positioned along potential reinvasion routes. This interpretation is consistent with studies showing that suitability surfaces often do not reliably predict landscape genetic patterns (Mateo-Sanchez et al. 2015; Milanesi et al. 2015; Sartor et al. 2022), while also aligning with tsetse work showing that integrating suitability and connectivity can reveal suitable but unconnected habitat patches that would be missed by habitat suitability alone (Bouyer et al. 2018).

#### Step 5. Post-hoc correlations with environmental predictors

Local Pearson correlations showed strong spatial heterogeneity in predictor-response relationships (Figure 6) that can be used to recover a more interpretable ecological picture of predicted connectivity (Balkenhol et al. 2009; Bishop et al. 2021; Pless et al. 2021; Vanhove and Launey 2023). Adding this post-hoc analysis allowed us to distinguish meaningful ecological differences between the North and South without requiring *a priori* knowledge of the potential mechanisms. The three broad patterns identified across predictors (Figure 6; Supplementary Figure S8), involving isothermality (BIO3), temperature variability (BIO4 and BIO7), and temperature and precipitation during the cooler and wetter periods (BIO11S and BIO13), all showed an approximate north-south transition. Together, these patterns suggest that climatic constraints on connectivity differ between Northern and Southern Uganda, potentially reflecting different balances of temperature-versus moisture-limitation.

Northern Uganda is generally flatter, drier, and has higher temperature seasonality and annual temperature range than in the South (Figure 6A), and the coldest season and wettest period occur during the wetter part of the year (Basalirwa 1995; Opiro et al. 2022). Under these conditions, the generally positive association of connectivity with BIO4 and BIO7 may indicate threshold effects of thermal variability on effective dispersal. At the same time, the generally negative association with BIO3, BIO11S, and BIO13 suggests that even modest increases in temperature or precipitation during the cooler and wetter part of the year may reduce connectivity, potentially pointing to thermal tolerance limits or dependence on cooler, humid refugia such as riverbanks and vegetated thickets.

Southern Uganda, in contrast, experiences a more buffered equatorial climate, with lower annual temperature range and more bimodal rainfall seasonality than the northern genetic cluster (Basalirwa 1995; Figure 6A), and the coldest season occurs during the relatively cool dry period from June through August, making the coldest season less extreme overall. Under these conditions, the generally negative association of connectivity with BIO4 and BIO7 may suggest that more stable thermal conditions facilitate connectivity across the landscape over multiple generations. Conversely, the generally positive association with BIO3, BIO11S, and BIO13 may indicate that warmer and wetter conditions reduce heat and desiccation stress, thereby supporting survival, greater fat reserves in emerging tenerals, and ultimately effective dispersal (Bursell 1958; Brightwell et al. 1992; Hargrove 2004; Kleynhans et al. 2014).

### Applications to vector control

In combination, connectivity and suitability can help define potential management units and identify regions where different vector control strategies may be most effective. Suitability identifies areas where *G. f. fuscipes* populations are likely to persist, whereas connectivity helps distinguish regions that may function as reinvasion corridors, areas that are more isolated and potentially more amenable to targeted suppression or local elimination, and areas where low suitability and low connectivity indicate comparatively low risk of both persistence and reinvasion. Our results provide specific examples of these patterns: highly connected habitat along the Albert Nile and northern shore of Lake Victoria may present substantial reinvasion risk, whereas isolated but suitable habitat near the Kenya-Uganda border north of Homa Bay and east of Lake Albert may represent potential targets for more localized interventions (Figure 4). These regional applications are considered in greater detail below.

The use of connectivity models to inform operational recommendations has precedent in tsetse control. In Senegal, stratified sampling, habitat suitability modeling, and population genetic evidence of isolation informed successful integrated control planning for G. palpalis gambiensis in the Niayes region (Solano et al. 2010; Dicko et al. 2014; Bouyer et al. 2015; Bouyer and Lancelot 2018; Ciss et al. 2019; Seck et al. 2024). Existing national control frameworks, including those used by the Coordinating Office for Control of Trypanosomiasis in Uganda (COCTU) and the Kenya Tsetse and Trypanosomiasis Eradication Council (KENTTEC), define tsetse management units and provide useful operational starting points, but may not capture within-belt variation in dispersal risk (Opiro et al. 2022; Bishop et al. 2021). Here, the North and South regions intersect broad tsetse control zones already recognized by COCTU and KENTTEC, and our results provide a more detailed interpretation of how regions differ in their potential to support persistence, reinvasion, or isolation.

In northern Uganda, the bivariate map highlights two regions with different implications for vector control. High-connectivity, high-suitability areas occur primarily in northwestern Uganda along the Albert Nile (pink; Figure 4). These regions are predicted to support persistent *G. f. fuscipes* populations with relatively high dispersal, suggesting that control efforts may be difficult to sustain unless treated areas are isolated from surrounding habitat or embedded within broader area-wide suppression efforts. This is particularly relevant along the Albert Nile, where eradication efforts should explicitly account for reinvasion risk. In contrast, high-connectivity, low-suitability areas are concentrated mainly around Lake Kyoga (yellow; Figure 4). These areas may not support large or stable year-round tsetse populations, but they could function as dispersal corridors that facilitate reinvasion or seasonal movement. This interpretation should be evaluated with additional field sampling and direct estimates of dispersal, especially because elevated modeled connectivity around Lake Kyoga could partly reflect admixture between northern and southern genetic clusters rather than current effective dispersal. Nonetheless, the model highlights Lake Kyoga as a region where existing policies aimed at limiting parasite and vector movement may warrant increased resources and targeted surveillance, especially during rainy seasons or periods of increased livestock movement. This is especially relevant in northern Uganda, where Opiro et al. (2022) identified the genetic transition zone between northwestern and northeastern Uganda as an epidemiologically important area between the two forms of HAT and emphasized the need for additional vector and parasite data to guide control interventions.

In southern Uganda and the Lake Victoria basin, the bivariate map identifies both connected and relatively isolated regions. High-connectivity, high-suitability areas along the northern shore of Lake Victoria are predicted to support persistent populations with relatively high dispersal, suggesting that local control may require broader coordination across connected habitat. In contrast, low-connectivity, high-suitability regions occur near the Kenya-Uganda border north of Homa Bay and in a patch east of Lake Albert (blue; Figure 4). These regions are predicted to support suitable habitat while remaining relatively isolated from surrounding populations, making them potential candidates for targeted suppression, local elimination, or field evaluation of novel AW-IPM tools, including SIT or emerging tools such as paratransgenesis (Solano et al. 2010; Aksoy et al. 2013; Vreysen et al. 2013; Dicko et al. 2014; Ciss et al. 2019; Seck et al. 2024; Hargrove et al. 2025). Although model predictions near range edges generally require cautious interpretation, these regions occur near known distributional limits, making edge artifacts less likely to explain the observed pattern.

More broadly, the use of landscape genetic connectivity to identify barriers, corridors, and relatively isolated populations has been applied across a wide range of taxa and management contexts (Balkenhol et al. 2009; Murphy et al. 2010; Shirk et al. 2017; Vanhove and Launey 2023). For vector management, the important general principle is that habitat suitability alone does not indicate whether a population is likely to remain isolated following intervention or be replenished through dispersal. Integrating suitability and connectivity therefore provides complementary information for identifying where localized interventions may be feasible and where coordinated management across connected habitat is more likely to be required.

### Strengths, limitations, and future directions in methodological approach

The main strength of this framework lies in its ability to predict relative connectivity and identify barriers, corridors, and relatively isolated populations, rather than to test specific mechanistic hypotheses or estimate globally uniform environmental effects. Recent work has emphasized that high predictive performance does not necessarily imply recovery of true causal resistance values when predictors are correlated or when modeled cost units diverge from the biological scale of genetic differentiation (Daniel et al. 2025). We therefore interpret the final surface primarily as a statistically calibrated prediction of relative connectivity rather than as a literal representation of pairwise CSE or environmental resistance. Consistent calibration across training sets and strong performance in both direct and least-cost-path-based LOPOCV increase confidence that the spatial patterns are robust (Figure 5), but do not establish the mechanisms responsible for those patterns.

Several methodological uncertainties remain. One concerns how best to interpret correlated predictors in random forest models. We used PCA-guided pruning to evaluate whether variable-importance rankings were robust to reduced predictor redundancy (Figure 3). Although the PCA-pruned comparison model performed nearly as well as the full model, we retained the complete mean-based predictor set for the final projected surface because our primary goal was to maximize spatial predictive performance rather than identify the smallest set of independent environmental predictors. Removing correlated predictors can simplify interpretation of individual variable effects, but may also discard predictive information shared among environmental variables. We therefore retained the full predictor set for spatial prediction while using the PCA-pruned comparison to evaluate the robustness of variable-importance rankings. If the primary goal were instead to identify a minimal set of environmental drivers or test specific hypotheses about their effects, a more strongly reduced predictor set and an inferential modeling framework designed around those hypotheses would be preferable.

The role of sampling density is also uncertain. Sampling density was included to account for uneven site coverage and retained in the distance-only baseline model for consistency across model formulations. During projection, this layer was neutralized to reduce direct imprinting of the sampling design on the final surface. However, its contribution is not straightforward to interpret, and future work should evaluate its effects more explicitly through targeted sensitivity analyses. More broadly, future methodological development should continue to examine how predictor scale, temporal mismatch, sampling structure, and alternative spatial validation schemes influence inference in applied landscape genetic models (Beninde et al. 2024; Daniel et al. 2025).

A further limitation is the use of microsatellite markers, which provide a relatively small set of predominantly neutral loci compared with genome-wide SNP data. Microsatellites are well suited for characterizing broad patterns of population structure and gene flow, but provide limited resolution for distinguishing landscape effects on neutral connectivity from environmentally associated genomic differentiation. Future applications using genome-wide SNP data could separately evaluate neutral and environmentally associated loci, allowing the framework to distinguish landscape features associated primarily with dispersal and gene flow from those potentially associated with local adaptation. Such an approach could provide a more complete picture of how both neutral and potentially adaptive processes contribute to spatial genetic structure and could identify environmental barriers relevant over different evolutionary timescales.

An especially promising future direction is extending this framework to forecast changes in genetic connectivity under future climate conditions. The present model characterizes connectivity under environmental conditions associated with the sampling period, but future projections are available for the bioclimatic variables used here. The predictive, multivariate structure of the machine-learning framework is particularly well suited to this application because it does not require connectivity to be reduced *a priori* to a small number of independently acting environmental variables. Instead, the fitted model can incorporate information distributed across correlated environmental predictors and project those relationships onto corresponding future climate surfaces. This provides an opportunity to forecast how connectivity may shift under alternative climate scenarios and identify potential changes in dispersal corridors, barriers, reinvasion risk, and management-unit boundaries. Such projections could extend the framework from informing control under current conditions to anticipating where long-term control strategies may need to adapt as climate changes.

## Conclusion

Our results addressed three objectives for translating landscape genetic models of connectivity into spatially explicit information for vector control. First, environmental predictors improved prediction of genetic connectivity relative to a distance-only model, demonstrating that environmental heterogeneity contributes to connectivity beyond geographic distance alone. Second, variable-importance and post-hoc spatial analyses identified environmental predictors associated with connectivity and revealed substantial spatial heterogeneity in these relationships, including broad differences between the North and South. Third, integrating predicted genetic connectivity with habitat suitability distinguished areas likely to support persistent and connected populations, potential routes of recolonization, and relatively isolated populations that may be more amenable to targeted suppression or local elimination.

Importantly, predictive performance of this analytical pipeline was strong across multiple complementary evaluations, including LOPOCV, permutation tests, direct evaluation of the projected connectivity surface, and comparison with distance-only baselines. The flexible nature of this machine-learning landscape genetics pipeline represents a strong opportunity to improve our ability to model, predict, and ultimately forecast features of pest populations relevant to management. The pipeline presented here offers several advantages: it does not require *a priori* specification of how individual environmental variables affect dispersal, it produces predictions that can be evaluated at both held-out sites and across projected spatial surfaces, it demonstrates strong generalizability in LOPOCV, it substantially improves prediction over distance-only approaches, and it can be applied to alternative versions of the environmental variables on which the models were trained. This last feature provides a particularly promising opportunity to project connectivity under future climate scenarios and forecast potential changes in corridors, barriers, and reinvasion risk.

For *G. f. fuscipes*, this framework may help guide more durable, spatially explicit interventions for human and animal African trypanosomiases by identifying where control should account for reinvasion risk, where surveillance should be prioritized, and where relatively isolated populations may be appropriate targets for local elimination or field evaluation of new control tools. Translating these landscape-level predictions into operational recommendations represents an important next step toward connecting landscape genetic inference with on-the-ground vector control. More broadly, this approach may be useful for other insect systems where dispersal and reinvasion are central to effective pest management.

## Supplementary Information

Supplementary File S1 includes additional methods and results for the newly genotyped western Kenya samples, population genetic analyses, and isolation-by-distance analyses, together with supporting figures and tables for the genetic connectivity modeling. Supplementary Figures S1–S3 present population genetic structure and isolation-by-distance analyses; Figures S4–S6 evaluate random forest convergence, projection scale and lake resistance, and iterative least-cost-path updating; Figure S7 presents leave-one-point-out cross-validation performance and baseline and permutation comparisons; and Figure S8 presents spatial correlations between environmental predictors and predicted genetic connectivity. Supplementary Tables S1–S3 provide supporting sample and population genetic information, including results for newly genotyped western Kenya samples, genetic clustering across sampled sites, and site-level population genetic summary statistics. Supplementary Tables S4–S6 present model-development and sensitivity analyses evaluating alternative genetic response formulations, river kernel bandwidth, and methods for summarizing projected resistance surfaces.

## Data Archiving and Availability Statement

Data inputs used in this study will be made available through the Dryad Digital Repository, with the DOI to be added after manuscript acceptance. All code and analysis scripts are available through the project’s GitHub repository: https://github.com/saarman/uganda-tsetse-LG.

## Acknowledgements

This work was supported indirectly by funding from the National Institutes of Health and the Fogarty International Center through grants supporting research and training on tsetse-transmitted African trypanosomiasis, tsetse evolutionary genetics, tsetse symbionts, trypanosome transmission biology, Spiroplasma effects on tsetse flies, parasitology and vector biology training, emergence of sleeping sickness foci in Uganda, and control of tsetse fly-transmitted diseases in Kenya (D43 TW007391, R01 AI068932, R01 AI139525, R01 AI158805, R21 AI163969, T32 AI007404, R01 AI051584, R03 TW008755, and U01 AI115648). This research also benefited from methodological discussions and brainstorming with Giuseppe Amatulli and Evlyn Pless, as well as field collections and research support provided by collaborators in Kenya and Uganda. The support and resources from the Center for High Performance Computing at the University of Utah are gratefully acknowledged.

