## Supplementary for "Mapping tsetse fly connectivity in Uganda with machine learning landscape genetics"

### Supplementary Methods

*New collections and microsatellite genotyping methods*

New samples were collected from six sites in western Kenya, selected because of their ecological proximity and geographic continuity with the Ugandan lakeshore sampling region. At each site, approximately six traps were placed at least 100 m apart and operated for 3 to 4 days. Four sites were collected in 2016 along the eastern shore of Lake Victoria: Budalangi (81-BUD), Bondo (82-BON), Suba (86-SUB), and Karungu (87-KAR). Two additional sites, Manga (84-MAN) and Kisasi (85-KIS), were originally sampled in 2009 and reported by Manangwa et al. (2017), but are included here through new genotyping and integration with the broader Uganda-western Kenya dataset. All flies were sampled using standardized field protocols coordinated by the Aksoy/Caccone tsetse research groups at Yale University. Biconical traps were used for collection, and flies were preserved individually in at least 80 percent ethanol. Metadata included sex, collection date, trap ID, and GPS coordinates.

In total, 166 individuals from the six Kenyan sites were genotyped, and 32 previously genotyped individuals from earlier studies were reanalyzed as validation controls. New samples were genotyped at 15 microsatellite loci using methods described in Echodu et al. (2011). PCR and fragment analysis conditions were kept consistent with earlier studies to ensure compatibility across datasets. DNA from a representative subset of individuals originally analyzed by Hyseni et al. (2012) was re-amplified and processed alongside the newly genotyped samples to confirm consistency in genotype scoring. Of the 15 loci, 11 overlapped with those used in prior studies and passed quality-control filters: A03b, B05, C07b, CAG29, GPCAG133, D05, D101, Gmm8, GmmB20b, GpC10, and Pgp28. New and existing datasets were then merged to generate a final dataset of 2,736 individuals genotyped at 11 loci.

*Population genetic clustering and summary statistics methods*

We evaluated spatial genetic structure using Bayesian and multivariate clustering approaches. Bayesian clustering was performed in STRUCTURE v2.3.4 using the admixture model with correlated allele frequencies (Pritchard et al. 2000). Ten replicate runs were conducted for each K from 1 to 20, with a burn-in of 50,000 iterations followed by 500,000 MCMC iterations. The optimal number of clusters was evaluated using the Evanno method in STRUCTURE Harvester (Evanno et al. 2005; Earl and vonHoldt 2012), and replicate runs were summarized using CLUMPAK (Kopelman et al. 2015). We also performed principal components analysis (PCA) using adegenet v2.1.4 (Jombart 2008), which does not assume Hardy-Weinberg or linkage equilibrium. Cluster membership for downstream analyses was determined based on concordant patterns from STRUCTURE and PCA.

Population genetic summary statistics were calculated by sampling site from the filtered microsatellite dataset using pegas v1.3 (Paradis 2010) for rarefied allelic richness, hierfstat v0.5.11 (Goudet and Jombart 2022) for FIS, observed heterozygosity (Ho), and expected heterozygosity (He), and poppr v2.9.8 (Kamvar et al. 2015) for the standardized index of association (rbarD).

*Isolation-by-distance and Mantel correlogram analysis methods*

Isolation by distance within each major cluster was assessed by comparing pairwise CSE with pairwise geographic distance among retained sites. Pairwise geographic distance was calculated from site coordinates in kilometers using haversine distance. Correlation between genetic and geographic distance within each cluster was tested using Mantel tests implemented with mantel.rtest in ade4 v1.7-23 (Mantel 1967; Dray and Dufour 2007), with 999 permutations. To evaluate the spatial scale of genetic structure, we constructed Mantel correlograms separately for the North and South clusters using mantel.correlog in vegan v2.7-1 (Oksanen et al. 2025). Correlograms were constructed using geographic distance classes defined by breakpoints at 0, 5, 10, 20, 40, 50, 100, 200, 300, 400, and 500 km, with 999 permutations and Holm correction for multiple testing.

### Supplementary Results

*New collections and microsatellite genotyping results*

A total of 166 individuals from six western Kenyan sites were genotyped at 15 microsatellite loci. Eleven loci overlapped with previously published datasets and were retained for integration into the full 2,736-individual dataset used in subsequent analyses. Among the newly genotyped populations, mean observed heterozygosity ranged from 0.261 to 0.537, mean expected heterozygosity ranged from 0.276 to 0.525, and mean number of alleles per locus ranged from 2.786 to 4.000 (Supplementary Table S1). Genetic diversity was generally higher at the northern sites, Busia and Siaya, and lower at Homa Bay and Migori. FIS ranged from -0.017 to 0.252, with significant deviations from zero at Busia (FIS = 0.078, $p = 0.032$) and Migori (FIS = 0.252, $p < 0.001$; Supplementary Table S1), while the remaining populations showed no significant deviations. Comparable statistics for previously published samples are available in Beadell et al. (2010), Hyseni et al. (2012), Kato et al. (2015), and Manangwa et al. (2017).

*Population genetic clustering and summary statistics results*

STRUCTURE and PCA supported two major genetic clusters for downstream modeling, corresponding to North and South (Figure 1). The Evanno method showed strongest support for $K = 2$ (Supplementary Figure S1), whereas PCA and STRUCTURE indicated additional substructure within the south cluster, including a distinct western subcluster. Sites 41 and 42 showed admixed ancestry and occupied intermediate positions in PCA space, while site 50 was greatly geographically isolated from the main sampling network and was removed because it did not have any neighboring sites within 50 km. The six newly genotyped western Kenya sites clustered with the South cluster, improving spatial coverage in the Lake Victoria region (Figure 1; Supplementary Table S2).

Summary statistics for all retained sites and the two major genetic clusters are reported in Table 1, with site-level values in Supplementary Table S3. Across the filtered dataset, site sample size ranged from 15 to 50 individuals and averaged 30.27. North had higher genetic diversity than south, with greater mean rarefied allelic richness (3.81 vs. 2.93) and mean expected heterozygosity (0.61 vs. 0.47). Pairwise CSE was slightly lower in the North than in the South (0.37 vs. 0.42).

*Isolation-by-distance and Mantel correlogram analysis results*

Mantel tests indicated strong positive isolation by distance within both major clusters, supporting inclusion of geographic distance as a predictor in downstream modeling (Supplementary Figure S2). Mantel correlation was slightly stronger in the North cluster ($r = 0.802$, $p = 0.001$) than in the South cluster ($r = 0.754$, $p = 0.001$). Mantel correlograms showed positive correlations at short to intermediate distance classes in both clusters, persisting through approximately 100 km (Supplementary Figure S3). Correlations declined toward zero and became negative at broader geographic scales, with the largest distance classes significantly negative in both clusters. Mantel correlograms suggest stable spatial structure within major genetic clusters over evolutionary time scales. Correlogram results indicating positive IBD until scales over 100 km suggest that isolation by distance has also been stable within genetic clusters over long evolutionary time scales. This further supports growing evidence that *G. f. fuscipes* displays long-term mutation-drift equilibrium within major genetic clusters in the North and South Uganda (Opiro et al. 2017; Saarman et al. 2019).

**Supplementary Figure S1.** STRUCTURE delta K plot based on the Evanno method. The strongest support was for K = 2, with additional substructure apparent at K = 3. Analyses used the admixture model with correlated allele frequencies in STRUCTURE v2.3.4, with 10 replicate runs per K, 50,000 burn-in iterations, and 500,000 MCMC iterations, summarized in CLUMPAK. Together with PCA and DAPC, these results supported the two major clusters (North and South) used in downstream analyses, while also indicating finer substructure among the North, West, and South groups.

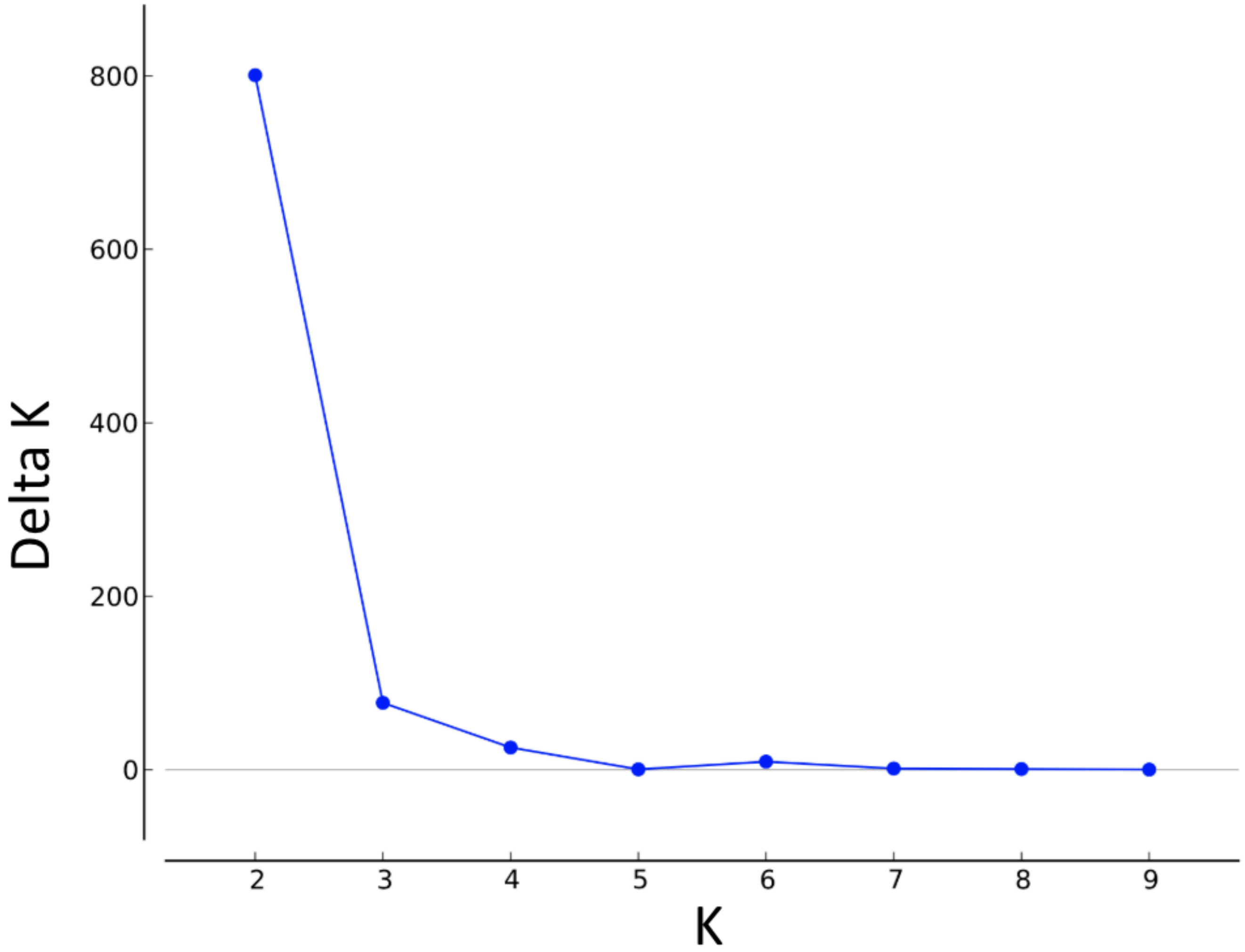

**Supplementary Figure S2.** Mantel tests and pairwise relationships between genetic and geographic distance within the two major genetic clusters retained for modeling. (a) Null distribution of Mantel correlation coefficients for the North cluster based on 999 permutations, with the observed Mantel correlation indicated by the black diamond. (b) Relationship between pairwise geographic distance (km) and Cavalli-Sforza and Edwards chord distance (CSE) for sites in the North cluster, with the fitted linear trend shown in red. (c) Null distribution of Mantel correlation coefficients for the South cluster based on 999 permutations, with the observed Mantel correlation indicated by the black diamond. (d) Relationship between pairwise geographic distance (km) and CSE for sites in the South cluster, with the fitted linear trend shown in red. Mantel tests indicated significant positive associations between genetic and geographic distance in both clusters (North: $r = 0.80$, $p < 0.001$; South: $r = 0.75$, $p < 0.001$).

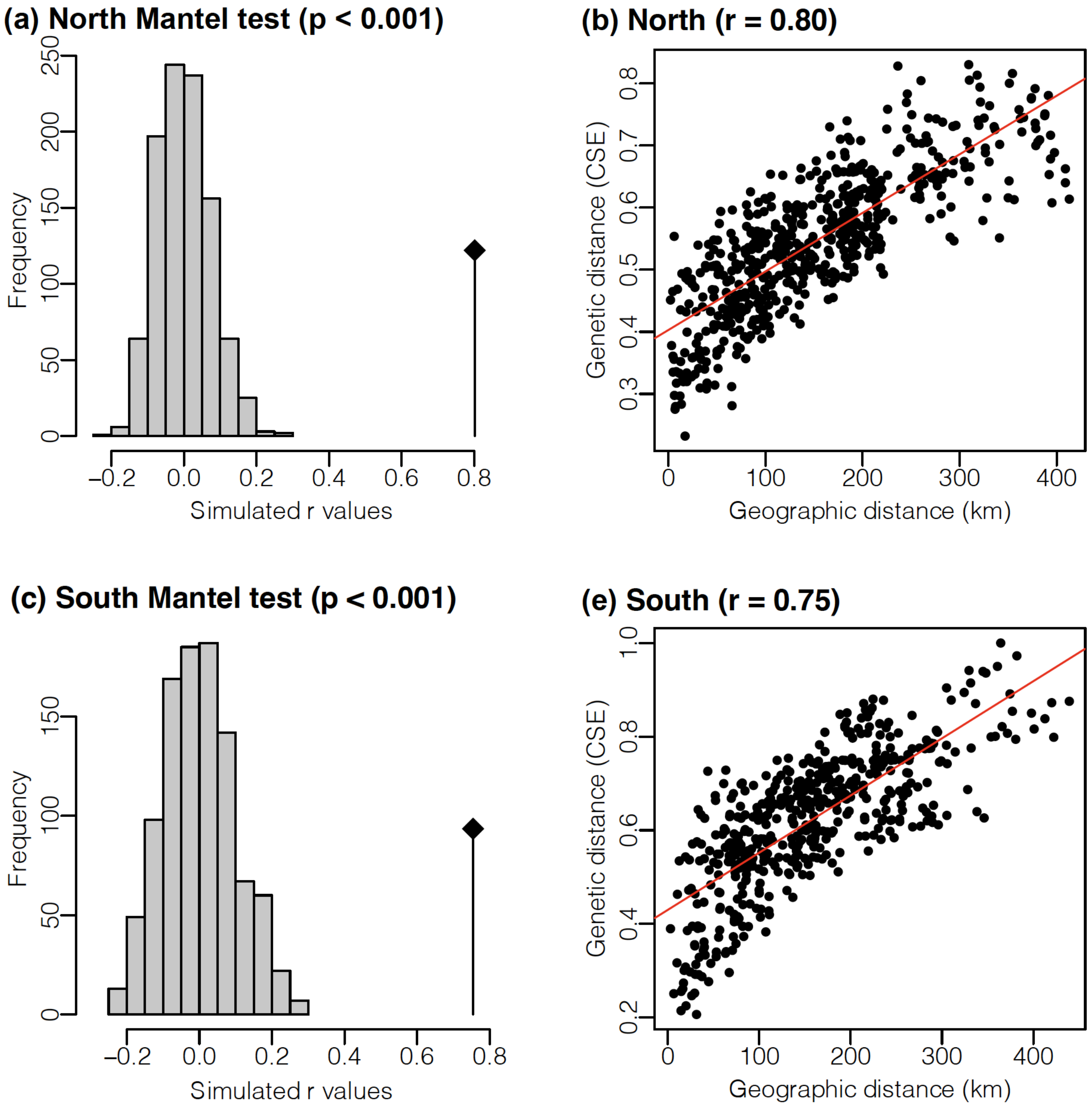

**Supplementary Figure S3.** Mantel correlograms for the two major genetic clusters retained for modeling. Panels show correlogram results for the (a) North and (b) South clusters, based on Mantel correlations between pairwise Cavalli-Sforza and Edwards chord distance (CSE) and pairwise geographic distance within successive distance classes. In both clusters, Mantel correlation was positive across short to intermediate distance classes, remained positive through approximately 100 km, and became negative at the broadest distance classes. Filled symbols indicate significant distance classes after Holm correction; open symbols indicate non-significant classes. The red horizontal line marks zero correlation.

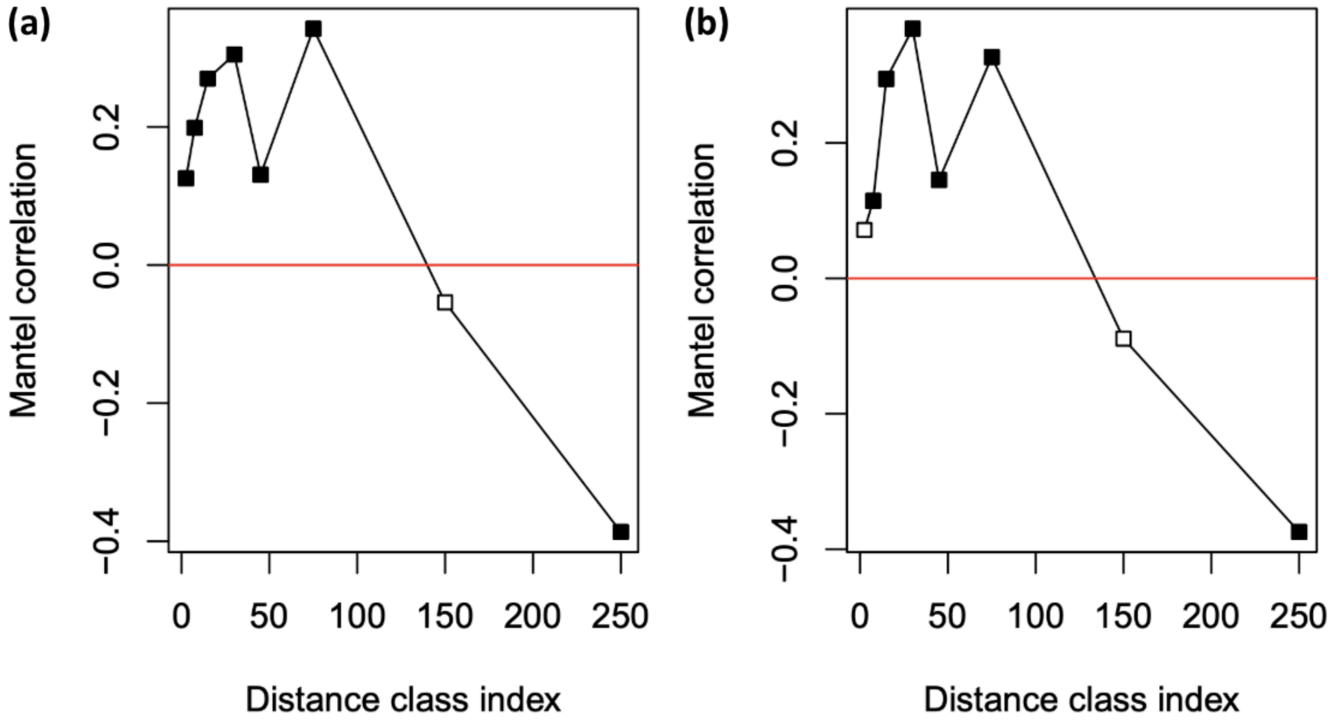

**Supplementary Figure S4.** Random forest convergence with increasing number of trees. Out-of-bag mean square error (OOB MSE) declined rapidly during initial model fitting and stabilized well before 500 trees, supporting the use of 500 trees in the final random forest model.

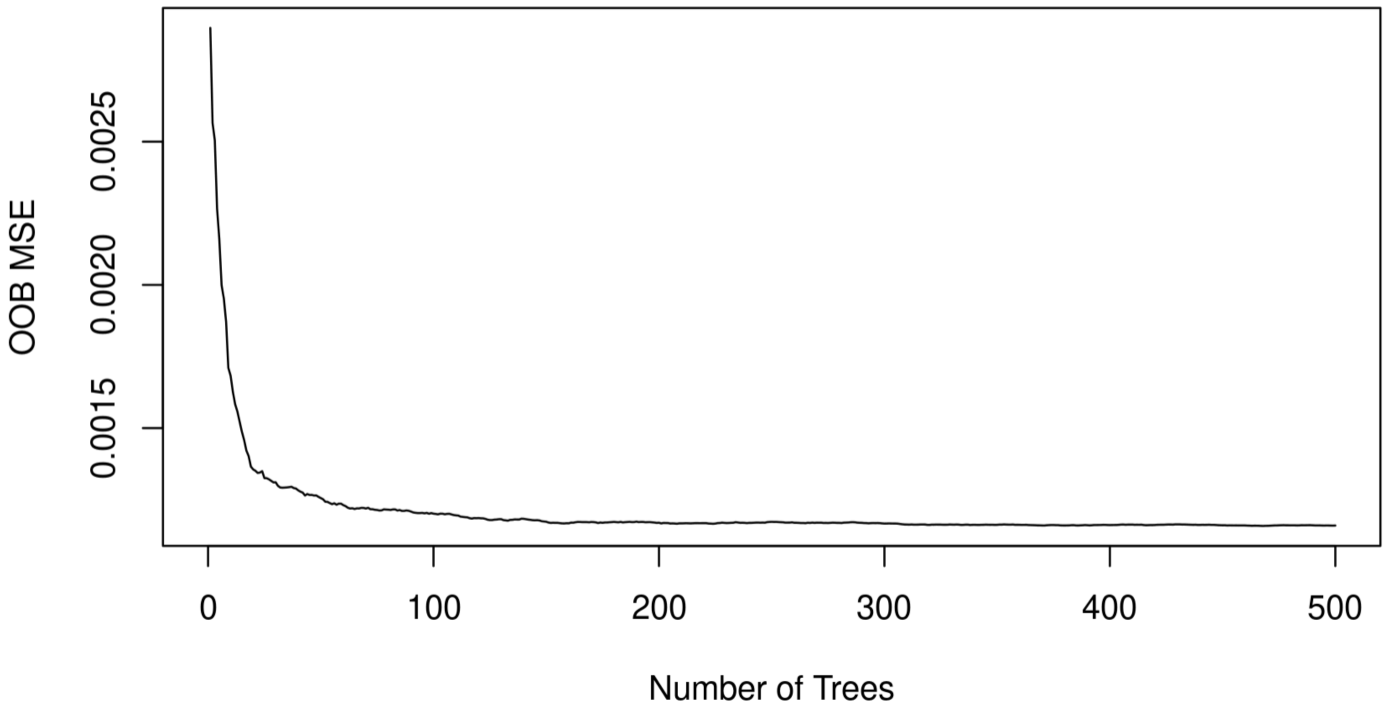

**Supplementary Figure S5.** Sensitivity of spatial evaluation of the genetic connectivity model to projection scale and lake resistance. Spatial performance was evaluated by comparing observed CSE genetic distance with calibrated predicted connectivity summarized along least-cost paths (LCP_sum), using spatial $R^2$, mean square error (MSE), and Spearman's r. Panels (a-c) show performance across projection scales from 1 to 100 km: (a) spatial $R^2$, (b) spatial MSE, and (c) Spearman's r. Performance declined slightly with increasing projection scale, supporting retention of the 1-km projection scale. Panels (d-f) show performance when lake resistance was set to 0.5, 1, 1.5, or 2 times the maximum predicted terrestrial resistance: (d) spatial $R^2$, (e) spatial MSE, and (f) Spearman's r. Performance was substantially lower at a multiplier of 0.5 and improved only modestly above the multiplier of 1 used in the final analysis, indicating that spatial evaluation was robust to equal or greater lake resistance.

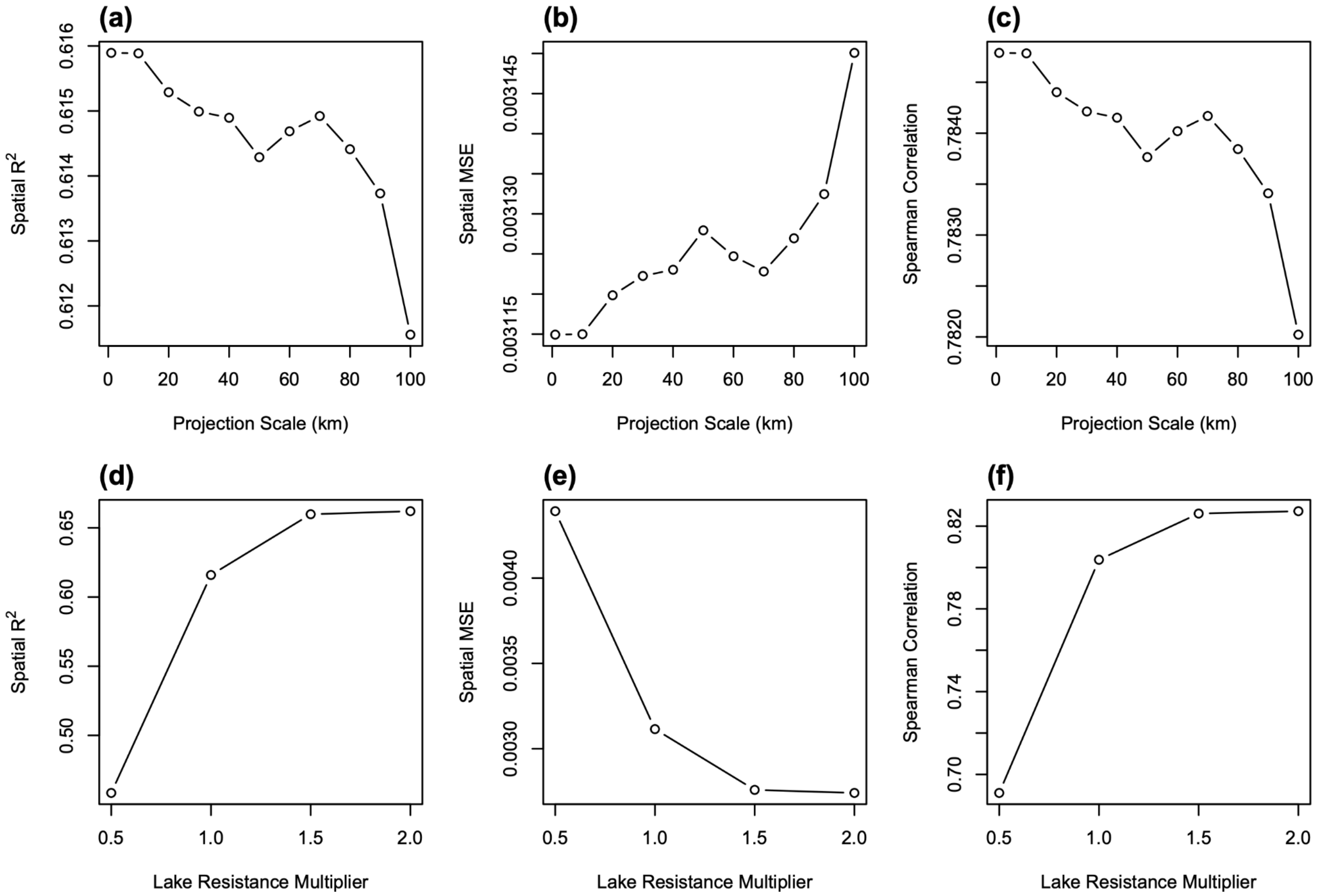

**Supplementary Figure S6.** Evaluation of iterative least-cost-path (LCP) model updating. Panels (a-b) show random forest model performance across 10 iterations based on direct model outputs: (a) percent variance explained and (b) out-of-bag mean square error (OOB MSE). Panel (c) shows mean absolute raster change between successive projected resistance surfaces. After an initial adjustment, subsequent iterations introduced only minor fluctuations in random forest performance and relatively small changes in the projected resistance surface, indicating that iterative updating did not consistently improve or substantially alter the model.

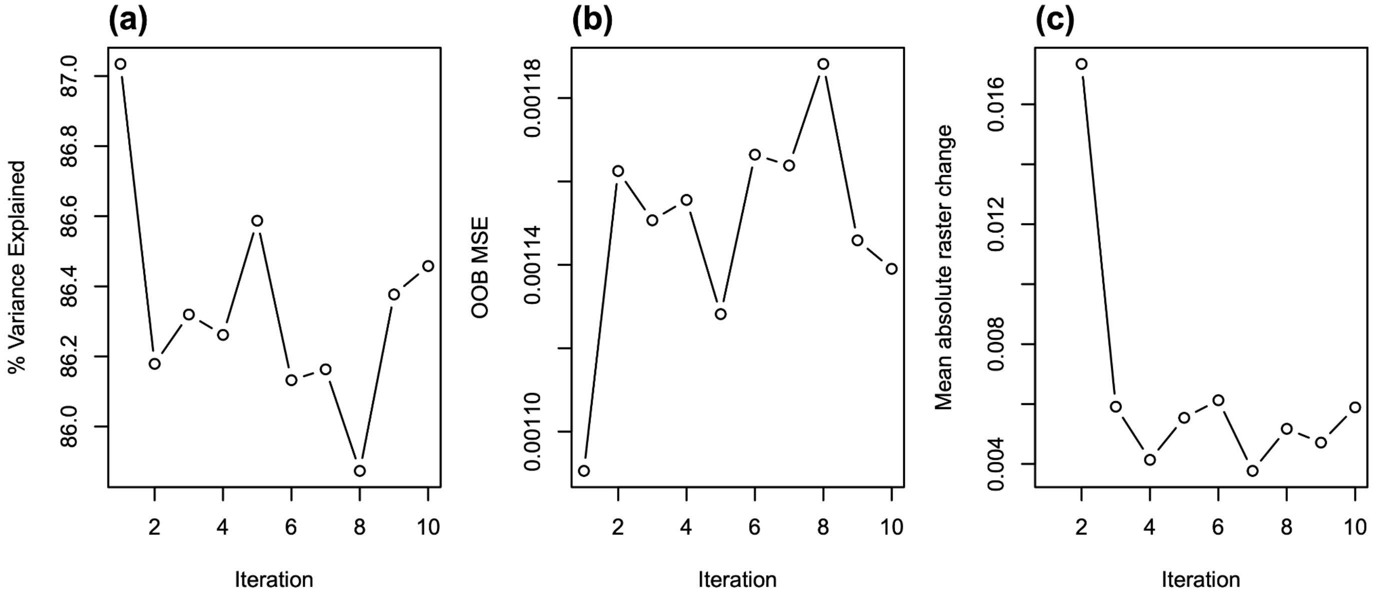

**Supplementary Figure S7.** Leave-one-point-out cross-validation (LOPOCV) performance of the genetic connectivity models and baseline comparisons. (a) Site-specific LOPOCV values from the random forest (RF) model mapped across held-out sites. (b) Site-specific LOPOCV values from the least-cost path (LCP) model mapped across held-out sites. In both maps, point size is proportional to fold-level Spearman's r, and point color indicates the genetic cluster used for regional model fitting, with North in blue and South in orange. (c) Distributions of fold-level Spearman's r values for the RF model, LCP model, baseline geographic-distance-only (GeoDist), and null-permutation results, shown separately for the North and South genetic clusters.

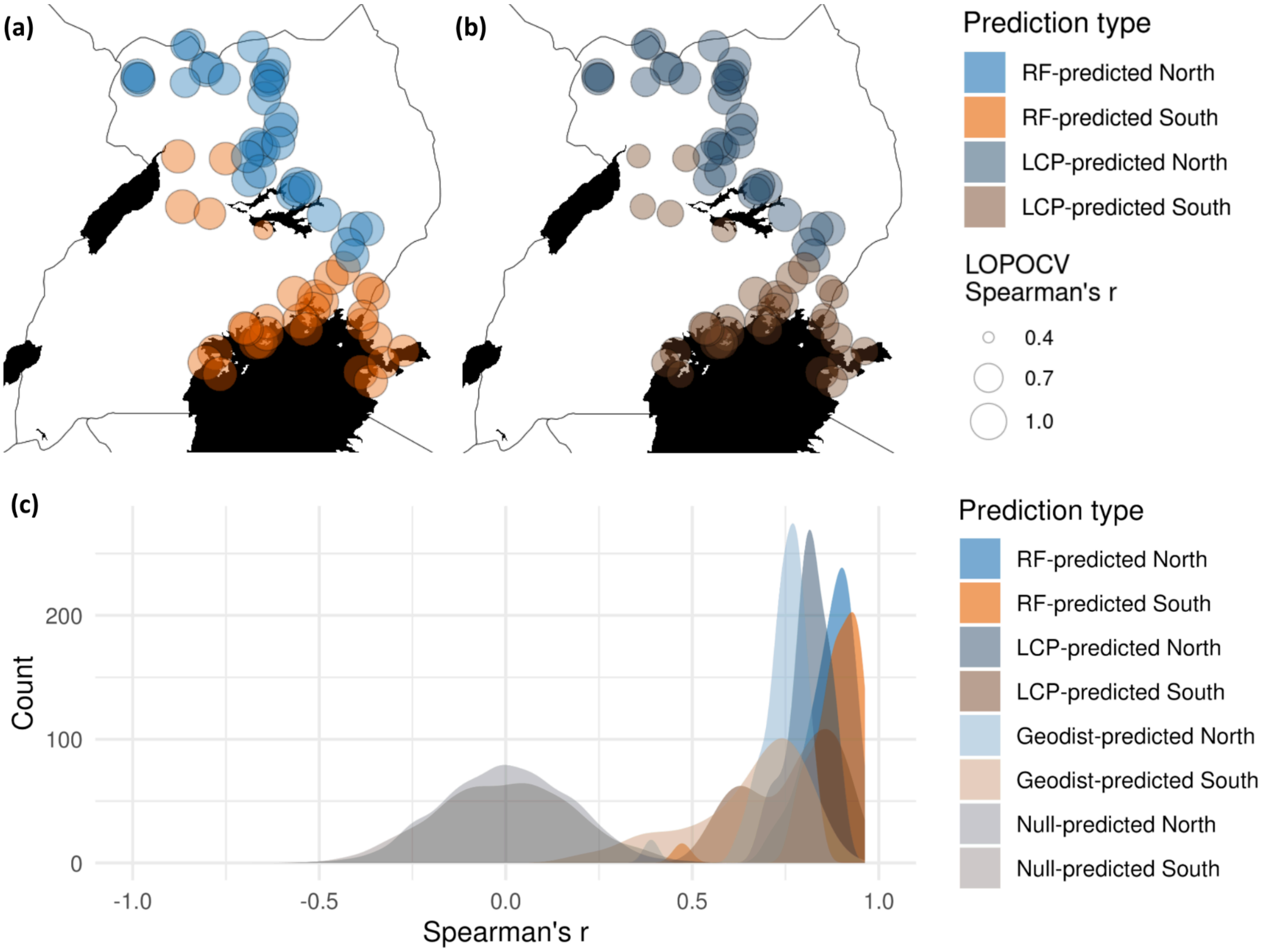

**Supplementary Figure S8.** Maps showing spatial variation in environmental associations with predicted genetic connectivity. (a) Raw values for five high-importance, relatively less-correlated environmental predictors: isothermality (BIO3), temperature seasonality (BIO4), temperature annual range (BIO7), mean temperature of the coldest season (BIO11S), and precipitation of the wettest month (BIO13). (b) Local Pearson correlations between each predictor and predicted genetic connectivity, calculated using a 21-pixel sliding window. Positive and negative values indicate corresponding local associations with connectivity. White areas indicate habitat suitability <0.05 and were masked from analysis.

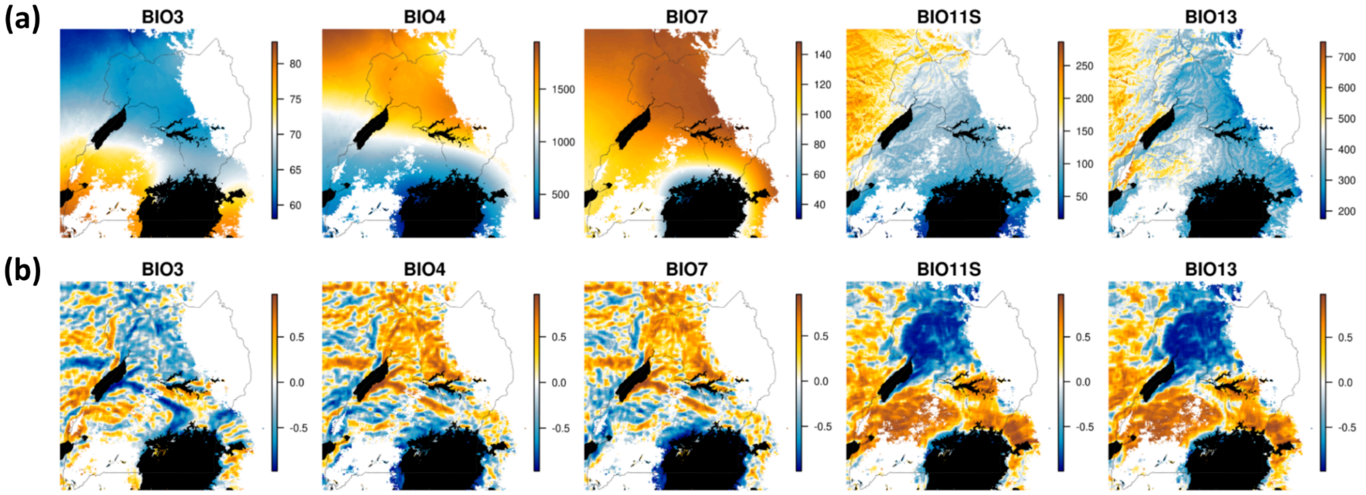

**Supplementary Table S1.** Population genetic analysis of newly genotyped samples from western Kenya. Sample site information and summary statistics for 166 Glossina fuscipes fuscipes individuals from six sites genotyped at 15 microsatellite loci. Eleven of these loci overlapped with previously published datasets and were retained for integration into subsequent population structure and landscape genetic analyses. N = number of individuals; HO = mean observed heterozygosity; HE = mean expected heterozygosity; NA = mean number of alleles across loci; FIS = inbreeding coefficient; FIS p-value tests whether FIS deviates significantly from zero. HO, HE, NA, FIS, and associated p-values were calculated using GenAlEx v6.41. Latitude and longitude are reported in decimal degrees. Comparable summary statistics for previously published samples are available in Beadell et al. (2010), Hyseni et al. (2012), Kato et al. (2015), and Manangwa et al. (2017).

| Site Code | Site Name | Lat. | Long. | N | Mean HO | Mean HE | Mean NA | $F_{IS}$ | p-value |
|---|---|---|---|---|---|---|---|---|---|
| 81-BUD | Busia | 0.1312 | 33.9845 | 24 | 0.461 | 0.488 | 4.000 | 0.0780 | 0.0318 |
| 82-BON | Siaya | -0.0466 | 34.2502 | 24 | 0.537 | 0.525 | 3.500 | 0.0010 | 0.4686 |
| 84-MAN | Manga | -0.3553 | 34.2513 | 35 | 0.395 | 0.382 | 3.571 | 0.0780 | 0.0882 |
| 85-KIS | Kisasi | -0.4764 | 33.9627 | 35 | 0.365 | 0.377 | 3.357 | -0.0170 | 0.3404 |
| 86-SUB | Homa Bay | -0.5930 | 34.0919 | 24 | 0.261 | 0.276 | 2.857 | 0.0460 | 0.1186 |
| 87-KAR | Migori | -0.8040 | 34.1088 | 24 | 0.286 | 0.372 | 2.786 | 0.2520 | 0.0002 |

**Supplementary Table S2.** Sampling site summary and genetic clustering results for *Glossina fuscipes fuscipes*. For each site, we report the site code (marked* if used in downstream modeling; marked ** if used in downstream modeling and represents a newly genotyped population), site name (N used if reduced), number of individuals genotyped (N), latitude, longitude, final cluster assignment (North, South, West, or Admixed), proportion of individuals assigned to the majority cluster by STRUCTURE (STR) for both K=3 and K=2, and data source (new or previously published). Anywhere that sample sizes were greater than 50, a random subset was selected for the genetic connectivity modeling.

| Site Code | Site Name | N | Lat. | Long. | Cluster | Sub-cluster | STR K=3 | STR K=2 | Data Source |
|---|---|---|---|---|---|---|---|---|---|
| 01-AIN* | Aina | 19 | 3.3044 | 31.1191 | North | North | 0.95 | 0.98 | Opiro 2017 |
| 02-GAN* | Gangu | 20 | 3.2521 | 31.1231 | North | North | 0.90 | 0.98 | Opiro 2017 |
| 03-DUK* | Duku | 25 | 3.2671 | 31.1352 | North | North | 0.92 | 0.96 | Opiro 2017 |
| 04-OM | Omugo | 15 | 3.2675 | 31.1431 | North | North | 1.00 | 0.98 | Beadell 2010; Opiro 2017 |
| 05-BLA | Belameling | 10 | 3.4790 | 31.5937 | North | North | 1.00 | 0.99 | Opiro 2017 |
| 06-LEA | Lea | 8 | 3.5923 | 31.6065 | North | North | 0.88 | 0.93 | Opiro 2017 |
| 07-OSG* | Osugo | 20 | 3.2110 | 31.7253 | North | North | 0.85 | 0.95 | Opiro 2017 |
| 08-MY* | Moyo | 15 | 3.6828 | 31.7270 | North | North | 0.87 | 0.96 | Opiro 2017 |
| 09-ORB* | Orubakulemi | 20 | 3.6922 | 31.7799 | North | North | 0.80 | 0.95 | Opiro 2017 |
| 10-PAG* | Pagirinya | 20 | 3.3777 | 31.9940 | North | North | 0.90 | 0.95 | Opiro 2017 |
| 11-OYA | Oringya | 9 | 3.4861 | 32.0105 | North | North | 0.89 | 0.97 | Opiro 2017 |
| 12-OLO* | Olobo | 24 | 3.4016 | 32.0110 | North | North | 0.96 | 0.98 | Opiro 2017 |
| 13-GOR | Gorodona | 25 | 3.2660 | 32.2081 | North | North | 1.00 | 0.98 | Opiro 2017 |
| 14-OKS* | Okidi | 26 | 3.2599 | 32.2241 | North | North | 0.88 | 0.98 | Opiro 2017 |
| 15-NGO* | Ngomoromo | 25 | 3.6687 | 32.5914 | North | North | 1.00 | 0.98 | Opiro 2017 |
| 16-PAW | Pawor | 13 | 3.6119 | 32.6819 | North | North | 1.00 | 0.98 | Opiro 2017 |
| 17-LAG* | Lagwel | 17 | 3.4407 | 32.8529 | North | North | 1.00 | 0.99 | Opiro 2017 |
| 18-BOL* | Bola | 25 | 3.2934 | 32.7822 | North | North | 0.96 | 0.98 | Opiro 2017 |
| 19-KTC* | Kitgum | 20 | 3.2823 | 32.8232 | North | North | 0.95 | 0.97 | Opiro 2017 |
| 20-TUM* | Tumangu | 20 | 3.2421 | 32.7608 | North | North | 0.95 | 0.97 | Opiro 2017 |
| 21-KT* | Kitgum | 17 | 3.1709 | 32.8049 | North | North | 0.76 | 0.92 | Beadell 2010 |
| 22-OMI* | Omido | 15 | 3.0113 | 32.7323 | North | North | 0.93 | 0.96 | Opiro 2017 |
| 23-PD | Pader | 13 | 3.0500 | 33.2167 | North | North | 0.69 | 0.91 | Opiro 2017 |
| 24-KIL* | Kilak | 21 | 2.7403 | 32.9502 | North | North | 0.95 | 0.98 | Opiro 2017 |
| 25-CHU* | Chua | 25 | 2.6069 | 32.9379 | North | North | 0.88 | 0.97 | Opiro 2017 |
| 26-OG* | Apala/Ogur | 37 | 2.4384 | 32.9124 | North | North | 0.92 | 0.98 | Beadell 2010 |
| 27-OCA* | Ocala | 20 | 2.4272 | 32.6285 | North | North | 0.95 | 0.98 | Opiro 2017 |
| 28-AKA* | Akayo-debe | 26 | 2.3722 | 32.6757 | North | North | 0.85 | 0.97 | Opiro 2017 |
| 29-KO | Kole | 15 | 2.3600 | 32.7152 | North | North | 0.60 | 0.95 | Opiro 2017 |
| 30-OLE* | Olepo | 24 | 2.3555 | 32.7159 | North | North | 1.00 | 0.99 | Opiro 2017 |
| 31-ACA* | Acanikoma | 25 | 2.2699 | 32.5209 | North | North | 0.72 | 0.94 | Opiro 2017 |
| 32-APU* | Aputu-Lwaa | 29 | 2.0794 | 32.6762 | North | North | 0.93 | 0.97 | Opiro 2017 |
| 33-AP* | Apac | 15 | 1.9756 | 32.5386 | North | North | 1.00 | 0.98 | Beadell 2010; Opiro 2017 |
| 34-AMI | Aminakwach | 25 | 1.9251 | 33.1562 | North | North | 1.00 | 0.99 | Echodu 2013; Opiro 2017 |
| 35-DK | Dokolo | 16 | 1.9166 | 33.1584 | North | North | 1.00 | 0.99 | Beadell 2010 |

| | | | | | | | | | |
|---|---|---|---|---|---|---|---|---|---|
| 36-UGT* | Kaber'ido | 64 | 1.9081 | 33.1602 | North | North | 0.98 | 0.99 | Opiro 2017 |
| 37-OT* | Otuboi | 40 | 1.8767 | 33.2557 | North | North | 0.98 | 0.99 | Echodu 2013 |
| 38-OCU* | Oculoi | 25 | 1.8470 | 33.1526 | North | North | 1.00 | 0.99 | Opiro 2017 |
| 39-OC | Oculoi | 20 | 1.8468 | 33.1539 | North | North | 0.95 | 0.99 | Echodu 2013 |
| 40-KAG* | Kangai | 20 | 1.8025 | 33.1034 | North | North | 0.95 | 0.98 | Opiro 2017 |
| 41-BUG | Bugonda | 18 | 1.6350 | 33.2900 | admix | admix | 0.83 | 0.66 | Burak 2018 |
| 42-BG | Bugondo | 13 | 1.6173 | 33.2982 | admix | admix | 0.77 | 0.76 | Beadell 2010 |
| 43-OS* | Osuguro | 32 | 1.5252 | 33.4970 | North | North | 0.88 | 0.96 | Beadell 2010 |
| 44-MK | Mukongoro | 24 | 1.3471 | 33.8933 | North | North | 1.00 | 0.99 | Echodu 2013 |
| 45-BKD* | Bukedea | 24 | 1.3491 | 34.0444 | North | North | 1.00 | 0.99 | Echodu 2013 |
| 46-PT* | Putiputi | 26 | 1.1488 | 33.7969 | North | North | 0.96 | 0.98 | Echodu 2013 |
| 47-BK* | Budaka | 40 | 1.0068 | 33.8574 | North | North | 0.90 | 0.96 | Echodu 2013 |
| 48-BN* | Bunghazi | 40 | 0.9355 | 33.9768 | North | North | 0.78 | 0.87 | Echodu 2013 |
| 50-KB | Kabunkanga | 40 | 0.9772 | 30.5465 | South | West | 0.95 | 0.74 | Beadell 2010 |
| 51-MF* | Murchison | 38 | 2.2720 | 31.6362 | South | West | 0.63 | 0.61 | Beadell 2010 |
| 52-KR* | Karuma | 55 | 2.2427 | 32.2397 | South | West | 0.65 | 0.57 | Echodu 2013 |
| 53-UWA | UWA | 25 | 2.2484 | 32.2491 | South | West | 0.60 | 0.61 | Opiro 2017 |
| 54-MS* | Masindi | 30 | 1.6303 | 31.6910 | South | West | 0.83 | 0.78 | Echodu 2013 |
| 55-KF* | Kafu | 32 | 1.5422 | 32.0406 | South | West | 0.96 | 0.81 | Echodu 2013, Opiro 2017 |
| 56-MA* | Masaka | 33 | -0.3564 | 31.9852 | South | South | 1.00 | 0.99 | Hyseni 2012 |
| 57-KG* | Kalangala Is. | 32 | -0.2267 | 32.1083 | South | South | 0.94 | 0.99 | Hyseni 2012 |
| 58-SS* | Ssese Is. | 40 | -0.5022 | 32.1691 | South | South | 0.98 | 0.99 | Hyseni 2012 |
| 59-EB* | Entebbe | 35 | 0.0823 | 32.4852 | South | South | 0.97 | 0.99 | Hyseni 2012 |
| 60-NA* | Nkumba | 33 | 0.0703 | 32.5085 | South | South | 1.00 | 0.99 | Beadell 2010 |
| 61-KO* | Koome Is. | 40 | -0.0911 | 32.6879 | South | South | 1.00 | 0.99 | Hyseni 2012 |
| 62-NS* | Nsazi Is. | 16 | 0.0127 | 32.7659 | South | South | 1.00 | 0.99 | Hyseni 2012 |
| 63-DB* | Damba Is. | 32 | 0.0127 | 32.7659 | South | South | 1.00 | 1.00 | Hyseni 2012 |
| 64-KL* | Kalengera | 40 | 0.1667 | 32.7667 | South | South | 0.98 | 0.99 | Beadell 2010 |
| 65-BZ* | Buziri Is. | 18 | 0.1716 | 33.1883 | South | South | 0.89 | 0.99 | Hyseni 2012 |
| 66-BY* | Bugaya Is | 62 | 0.0675 | 33.2684 | South | South | 0.92 | 0.98 | Hyseni 2012 |
| 67-WAM | Wambogo | 14 | 0.3687 | 33.3455 | South | South | 0.86 | 0.97 | Saarman 2018 |
| 68-LI* | Lingira Is. | 46 | 0.3168 | 33.3532 | South | South | 0.89 | 0.98 | Hyseni 2012 |
| 69-BV* | Buvuma | 60 | 0.4742 | 33.3824 | South | South | 0.93 | 0.99 | Hyseni 2012 |
| 70-MGG* | Mayuge | 59 | 0.4222 | 33.4559 | South | South | 0.95 | 0.98 | Echodu 2013 |
| 71-BD* | Budondo | 35 | 0.5208 | 33.1209 | South | South | 0.86 | 0.98 | Hyseni 2012 |
| 72-JN* | Junda | 40 | 1.3367 | 32.7247 | South | South | 0.60 | 0.98 | Echodu 2013 |
| 73-IGG* | Iganga | 65 | 0.7312 | 33.5861 | South | South | 0.95 | 0.99 | Echodu 2013 |
| 74-NAM* | Namutumba | 15 | 0.8450 | 33.7406 | South | South | 1.00 | 0.99 | Echodu 2013 |
| 75-KIS | Kisoko | 8 | 0.8276 | 34.1700 | South | South | 1.00 | 0.99 | Echodu 2013 |
| 76-TB* | Tuba | 28 | 0.5916 | 34.0586 | South | South | 0.82 | 0.91 | Echodu 2013 |
| 77-NB | Nambogo | 17 | 0.5918 | 34.0590 | South | South | 0.94 | 0.97 | Echodu 2013 |
| 78-OK* | Okame | 40 | 0.5194 | 34.1190 | South | South | 0.98 | 0.99 | Echodu 2013 |
| 79-BU* | Busiime | 40 | 0.2434 | 33.9932 | South | South | 0.97 | 0.99 | Beadell 2010; Echodu 2013 |
| 80-SA | Sangalo | 15 | 0.2420 | 33.9930 | South | South | 0.93 | 0.98 | Beadell 2010; Echodu 2013 |

| | | | | | | | | | |
|---|---|---|---|---|---|---|---|---|---|
| 81-BUD** | Busia | 24 | 0.1312 | 33.9845 | South | South | 1.00 | 0.99 | This study |
| 82-BON** | Siaya | 24 | -0.0466 | 34.1502 | South | South | 0.92 | 0.97 | This study |
| 83-ND* | Ndere KE | 40 | -0.2055 | 34.5095 | South | South | 1.00 | 0.99 | Beadell 2010 |
| 84-MAN** | Manga | 35 | -0.3553 | 34.2513 | South | South | 1.00 | 0.99 | This study |
| 85-KSS** | Kisasi | 35 | -0.4764 | 33.9627 | South | South | 0.97 | 0.98 | This study |
| 86-SUB** | Homo Bay | 24 | -0.5930 | 34.0919 | South | South | 1.00 | 0.99 | This study |
| 87-KAR** | Migori | 24 | -0.8040 | 34.1088 | South | South | 1.00 | 1.00 | This study |

**Supplementary Table S3.** Site-level summary statistics for all sampled populations, including sample size (N), allelic richness (AR), multilocus linkage disequilibrium (rbarD), inbreeding coefficient (FIS), observed heterozygosity (Ho), and expected heterozygosity (He). Summary rows report mean, minimum, and maximum values across all sites and within the North and South groups.

| Site Code | N | AR | rbarD | FIS | Ho | He |
|---|---|---|---|---|---|---|
| 01-AIN | 19 | 4.5832 | 0.0042 | 0.0410 | 0.6910 | 0.7210 |
| 02-GAN | 20 | 4.2607 | -0.0030 | 0.1890 | 0.5700 | 0.6980 |
| 03-DUK | 25 | 4.3249 | 0.0018 | 0.1550 | 0.5750 | 0.6880 |
| 07-OSG | 20 | 4.2983 | -0.0286 | 0.1370 | 0.5850 | 0.6890 |
| 08-MY | 15 | 4.0851 | 0.0007 | 0.0820 | 0.5760 | 0.6410 |
| 09-ORB | 20 | 4.1245 | -0.0064 | 0.1950 | 0.5320 | 0.6490 |
| 10-PAG | 20 | 3.9893 | 0.0003 | 0.1600 | 0.5500 | 0.6510 |
| 12-OLO | 24 | 3.7995 | 0.0013 | 0.0360 | 0.5870 | 0.6080 |
| 14-OKS | 26 | 4.1590 | -0.0089 | 0.0740 | 0.5620 | 0.6200 |
| 15-NGO | 25 | 4.0357 | -0.0066 | 0.0440 | 0.6200 | 0.6450 |
| 17-LAG | 17 | 3.6718 | 0.0680 | 0.1380 | 0.4880 | 0.5690 |
| 18-BOL | 25 | 3.9710 | 0.0062 | 0.0460 | 0.5820 | 0.6140 |
| 19-KTC | 20 | 4.0016 | 0.0173 | 0.0720 | 0.5750 | 0.6360 |
| 20-TUM | 20 | 3.9208 | 0.0124 | 0.0430 | 0.5640 | 0.6060 |
| 21-KT | 17 | 3.9542 | 0.0077 | 0.1260 | 0.5570 | 0.6370 |
| 22-OMI | 15 | 4.3067 | 0.0321 | 0.1550 | 0.5600 | 0.6350 |
| 24-KIL | 21 | 4.0539 | 0.0094 | 0.0480 | 0.6040 | 0.6410 |
| 25-CHU | 25 | 3.8828 | 0.0098 | 0.0030 | 0.5930 | 0.6010 |
| 26-OG | 37 | 4.0055 | -0.0111 | 0.0160 | 0.6230 | 0.6220 |
| 27-OCA | 20 | 4.1959 | -0.0091 | 0.0140 | 0.6410 | 0.6670 |
| 28-AKA | 26 | 3.9554 | 0.0066 | -0.0110 | 0.6210 | 0.6170 |
| 30-OLE | 24 | 3.7906 | 0.0113 | 0.0270 | 0.5770 | 0.5970 |
| 31-ACA | 25 | 4.0117 | 0.0208 | 0.0440 | 0.5690 | 0.6070 |
| 32-APU | 29 | 3.7365 | -0.0065 | 0.0680 | 0.5560 | 0.6030 |
| 33-AP | 15 | 3.8191 | 0.0353 | 0.0850 | 0.5560 | 0.6080 |
| 36-UGT | 50 | 3.6158 | 0.0091 | 0.0820 | 0.5360 | 0.5840 |
| 37-OT | 40 | 3.4914 | 0.0065 | 0.0500 | 0.5380 | 0.5700 |
| 38-OCU | 25 | 3.4153 | -0.0058 | -0.0030 | 0.5550 | 0.5690 |
| 40-KAG | 20 | 3.4985 | 0.0298 | 0.2040 | 0.4790 | 0.5740 |
| 43-OS | 32 | 3.3525 | 0.0013 | 0.0660 | 0.5140 | 0.5470 |
| 44-MK | 24 | 2.7319 | -0.0108 | -0.0640 | 0.4590 | 0.4320 |
| 45-BKD | 24 | 2.8992 | -0.0077 | 0.0680 | 0.4170 | 0.4530 |
| 46-PT | 26 | 2.9999 | -0.0020 | 0.0680 | 0.4840 | 0.5140 |
| 47-BK | 40 | 3.0986 | 0.0113 | 0.0870 | 0.4580 | 0.5030 |
| 48-BN | 40 | 3.4521 | 0.0320 | 0.0840 | 0.5240 | 0.5780 |
| 51-MF | 38 | 4.0476 | -0.0016 | 0.0170 | 0.6260 | 0.6350 |
| 52-KR | 50 | 4.1306 | -0.0008 | 0.1000 | 0.5910 | 0.6570 |
| 54-MS | 30 | 3.6076 | 0.0144 | 0.0530 | 0.5590 | 0.5830 |
| 55-KAF | 50 | 3.1582 | 0.0036 | 0.0760 | 0.4650 | 0.4970 |

| | | | | | | |
|---|---|---|---|---|---|---|
| 56-MA | 33 | 2.2973 | 0.0283 | -0.0950 | 0.3380 | 0.3040 |
| 57-KG | 32 | 2.2609 | -0.0005 | 0.0120 | 0.3100 | 0.3160 |
| 58-SS | 40 | 2.4636 | 0.0047 | 0.0350 | 0.3240 | 0.3430 |
| 59-EB | 35 | 2.9105 | 0.0488 | 0.0320 | 0.4660 | 0.4880 |
| 60-NA | 33 | 2.4638 | 0.0160 | -0.1040 | 0.5090 | 0.4670 |
| 61-KO | 40 | 2.4447 | -0.0087 | 0.0120 | 0.3770 | 0.3830 |
| 62-NS | 16 | 2.7151 | 0.0244 | 0.1030 | 0.3870 | 0.4080 |
| 63-DB | 32 | 2.4652 | -0.0055 | -0.0290 | 0.4390 | 0.4230 |
| 64-KL | 40 | 2.7423 | 0.0059 | 0.0080 | 0.4870 | 0.4780 |
| 65-BZ | 18 | 2.9723 | 0.0072 | 0.0700 | 0.4570 | 0.4860 |
| 66-BY | 50 | 3.2028 | 0.0095 | 0.0190 | 0.4930 | 0.5080 |
| 68-LI | 46 | 3.4096 | -0.0002 | -0.0120 | 0.5340 | 0.5240 |
| 69-BV | 50 | 3.2287 | -0.0029 | 0.0630 | 0.5020 | 0.5330 |
| 70-MGG | 50 | 3.3019 | -0.0060 | -0.0060 | 0.5190 | 0.5190 |
| 71-BD | 35 | 3.2033 | -0.0124 | 0.0150 | 0.5070 | 0.5100 |
| 72-JN | 40 | 2.7760 | 0.0074 | 0.0220 | 0.4350 | 0.4450 |
| 73-IGG | 50 | 3.2380 | 0.0048 | 0.0770 | 0.4900 | 0.5290 |
| 74-NAM | 15 | 3.4435 | -0.0195 | -0.0260 | 0.5590 | 0.5440 |
| 76-TB | 28 | 3.0454 | -0.0002 | 0.0250 | 0.4960 | 0.5120 |
| 78-OK | 50 | 3.0886 | -0.0104 | 0.0140 | 0.5050 | 0.5120 |
| 79-BU | 50 | 3.1273 | 0.0124 | 0.0940 | 0.4600 | 0.5040 |
| 81-BUD | 24 | 3.1243 | 0.0081 | 0.0700 | 0.4850 | 0.5130 |
| 82-BON | 24 | 3.0888 | 0.0440 | -0.0800 | 0.5770 | 0.5350 |
| 83-ND | 40 | 2.2515 | 0.0258 | 0.0070 | 0.3550 | 0.3550 |
| 84-MAN | 35 | 2.6285 | 0.0093 | -0.0070 | 0.4030 | 0.3950 |
| 85-KSS | 35 | 2.4333 | -0.0070 | 0.0470 | 0.3450 | 0.3640 |
| 86-SUB | 24 | 2.1081 | 0.0072 | 0.0270 | 0.2850 | 0.2920 |
| 87-KAR | 24 | 2.3100 | 0.0045 | 0.2970 | 0.2620 | 0.3810 |

**Supplementary Table S4. Comparison of candidate response formulations evaluated during model development.** Spatially evaluated spearman's correlation (cor.), $R^2$, root mean square error (RMSE), and mean absolute error (MAE) represent the predictive performance of the models of pairwise Cavalli-Sforza and Edwards (CSE) genetic distance after projection to the landscape and summed along least-cost paths. Three candidate response formulations were evaluated: (i) raw pairwise Cavalli-Sforza and Edwards (CSE) genetic distance, (ii) CSE residuals from a linear regression of CSE against Euclidean geographic distance (i.e., isolation-by-distance residuals), and (iii) CSE per kilometer (CSE/km), calculated by dividing pairwise CSE by the number of 1-km pixels traversed along the least-cost path. Raw CSE consistently produced the strongest predictive performance (highest correlation and $R^2$, lowest RMSE and MAE) and was therefore retained for all subsequent analyses.

| **Response formulation** | **Spatial cor.** | **Spatial $R^2$** | **Spatial RMSE** | **Spatial MAE** | **Final model** |
|---|---|---|---|---|---|
| CSE (raw genetic distance) | 0.785 | 0.616 | 0.056 | 0.045 | Yes |
| CSE isolation-by-distance residuals | -0.054 | 0.003 | 0.090 | 0.071 | No |
| CSE per kilometer (CSE/km) | 0.642 | 0.410 | 0.069 | 0.055 | No |

**Supplementary Table S5.** Sensitivity of genetic connectivity model performance to river kernel bandwidth. Random forest model performance was evaluated across river kernel-density bandwidths of 1, 2, 3, 5, and 10 km while holding all other predictors and model settings constant. Percent variance explained and out-of-bag mean square error (OOB MSE) were nearly identical across bandwidths, indicating that model performance was insensitive to this analytical choice.

| **River Kernel Bandwidth** | **% Var. Explained** | **OOB MSE** | **Biological interpretation** | **Final model** |
|---|---|---|---|---|
| 1 km | 85.82 | 0.00115 | Within lifetime dispersal scale; captures local riverine conditions relevant to pupal deposition. | No |
| 2 km | 85.79 | 0.00115 | Within lifetime dispersal scale; captures local riverine conditions relevant to pupal deposition. | No |
| 3 km | 85.81 | 0.00115 | At the upper end of the lifetime dispersal scale; captures local riverine conditions relevant to pupal deposition. | Yes |
| 5 km | 85.88 | 0.00114 | Extends beyond the typical lifetime dispersal scale; genetic differentiation has been detected at this scale. | No |
| 10 km | 85.74 | 0.00116 | Broad landscape scale substantially exceeds typical lifetime dispersal ability. | No |

**Supplementary Table S6. Comparison of candidate methods for summarizing projected resistance surfaces into pairwise estimates of genetic connectivity.** Spatially evaluated Spearman's correlation (cor.), $R^2$, root mean square error (RMSE), and mean absolute error (MAE) represent the predictive performance of the models of pairwise Cavalli-Sforza and Edwards (CSE) genetic distance after projection to the landscape and summarized along least-cost paths. Summarization methods included sum, mean, and a circuit-based effective resistance (CBER) procedure. Sum of the resistance values along least-cost paths (LCP-sum) produced the best predictive performance and was therefore retained for all subsequent analyses.

| **Spatial summarization method** | **Spatial cor.** | **Spatial $R^2$** | **Spatial RMSE** | **Spatial MAE** | **Final model** |
|---|---|---|---|---|---|
| Least-cost path resistance sum (LCP-sum) | 0.785 | 0.616 | 0.056 | 0.045 | Yes |
| Circuit-based effective resistance (CBER) | 0.567 | 0.560 | 0.059 | 0.0463 | No |
| Least-cost path mean resistance (LCP-mean) | 0.310 | 0.100 | 0.085 | 0.067 | No |
